# Genome reorganization and its functional impact during breast cancer progression

**DOI:** 10.1101/2025.05.14.654144

**Authors:** Kathleen S. Metz Reed, Andrew Fritz, Haley Greenyer, Kerstin Heselmeyer-Haddad, Seth Frietze, Janet Stein, Gary Stein, Tom Misteli

## Abstract

Cancer progression involves extensive alterations in epigenetic and gene expression programs, but the accompanying changes in higher-order genome organization remain less well understood. Using high-resolution Micro-C mapping in the MCF10 cell model of breast cancer, we profiled chromatin compartments, topologically associated domains, and chromatin loops. We find large-scale compartmental shifts occur predominantly in early stages of cancer development, with more fine-scale structural changes in TADs and looping accumulating during the later transition to metastasis. Relating these chromatin features to gene expression and enhancer-associated histone marks revealed that many differentially expressed genes are physically connected to distal regulatory elements. While enhancer-promoter contact frequency and distal enhancer activity correlated with gene expression, strong changes in chromatin looping were relatively infrequent during progression, suggesting that alterations in chromatin contacts are not globally necessary but may facilitate gene regulation at a subset of genes. These results elucidate the connection between gene regulation and genome remodeling in a cell-based cancer progression model.

## INTRODUCTION

The eukaryotic genome is highly organized in the cell nucleus. Amongst the most prominent structural features are kilobase-sized chromatin loops, medium-scale topologically associating domains (TADs) and higher-order compartments (1–5). How chromatin organization contributes to epigenetic control of gene regulation, including in physiological and pathological settings, such as cancer, remains only partially understood (6–9).

A prominent mechanism to generate higher order chromatin structures is loop extrusion (10–12). During this process, the multi-component cohesin complex is loaded onto chromatin and, using its intrinsic molecular motor activity, extrudes chromatin bidirectionally along the genome to form a loop or a domain until it encounters the major chromatin architectural protein CTCF bound to convergently oriented binding sites. The encounter of cohesin with bound CTCF stalls the extrusion process and generates a chromatin loop or TAD. While loop extrusion has universally been implicated in formation of chromatin loops and domains, formation of chromatin features by other mechanisms have also been observed, especially at a smaller scale (13–16).

A common property of higher order chromatin folding is that the resulting loops, domains and compartments, bring distal genome elements into spatial proximity and into proximity of regulatory elements to their target genes has been implicated in gene regulation (2). For example, it has been suggested that one function of TADs is to facilitate the interaction of regulatory elements, particularly gene enhancers, with their target genes located within the same TAD (17–21). Similarly, long-range interactions via chromatin loops are thought to be essential at some loci to bring gene enhancers into proximity to their target promoters (22–26). However, regulation of many gene loci also appears independent of chromatin organization, and enhancer-promoter proximity often does not correlate with gene activity (27–30). In fact, acute depletion of cohesin revealed genome-wide disruption of chromatin organization but surprisingly limited impact on gene expression (31). A possible explanation for these divergent findings is that chromatin organization may be functionally more relevant to bring about changes in gene activity rather than maintenance of gene expression as suggested by several cohesin depletion studies (30,32,33).

Genome organization is likely relevant for cancer and its progression. Mutations in loop extrusion machinery, such as cohesin and the cohesin processivity factor NIPBL or at CTCF binding sites have been reported in many cancers (6,7,34–36). In addition, many structural variants (SVs) such as deletions, duplications, and translocations, have been documented in various cancer subtypes where SVs and the ensuing reorganization around them can lead to aberrant gene regulation (37–39).

Despite these observations, the full extent of genome reorganization during cancer progression, and its functional consequences, remains largely unknown. Several studies primarily focused on large scale reorganizations have found changes in higher-order chromatin organization such as chromosome clustering and dynamic compartments, some of which correlated with changes in differentially expressed oncogenes and enhancers (40–48). Analysis of TADs in various cancers have found mixed results, with some studies pointing to increased TADs and gained boundaries and others observing more stable TAD organization or weakened boundaries (42,46,47,49,50). Furthermore, cancer-associated structural variants, such as chromosomal translocations, have been related to altered gene expression, for example via enhancer-hijacking (51,52). However, how local chromatin loops and TADs are restructured during oncogenic reprogramming and how these changes relate to cancer-associated gene expression has not been well documented.

We address the question of how local and global chromatin organization changes during cancer progression, and how they related to cancer gene expression programs, by generating high-resolution Micro-C maps in the well-established MCF10 breast cancer progression model (53). This cancer model consists of three epithelial cell lines that were all originally derived from the same non-cancerous patient (54). MCF10A is an adherent epithelial non-cancerous cell line that spontaneously immortalized from the initial patient sample. Pre-malignant MCF10AT1 cells were derived from MCF10A cells by overexpressing mutant Ha-Ras oncogene followed by long-term passage as mouse xenografts (55). In immunocompromised mice, MCF10AT1 cells form precancerous lesions, and approximately 25% progress to invasive carcinoma over time. The MCF10CA1a cell line is derived from metastatic tumors generated from xenografted MCF10AT1 cells and is considered fully malignant, metastatic, and highly aggressive, forming tumors in 100% of xenografted mice which can metastasize rapidly to the lung (56). The MCF10 progression series represents a spectrum of cells that share a similar genetic background but are increasingly more cancerous and they have been widely used to study the genetic and epigenetic changes that occur during cancer progression and epithelial-mesenchymal transitions (EMT) (41,57–65).

In this study, comparing fine-scale chromatin organization and other epigenetic features in the MCF10 cancer progression model has allowed us to identify changes in genome reorganization and relate them to changes in gene expression, including of cancer progression-associated genes. Our results provide novel insights into the principles of chromatin-mediated gene regulation and into the dynamic structure-function relationship contributing to genome regulation derived from an *in vitro* cancer progression model.

## RESULTS

### Mapping global and local genome organization across breast cancer progression

To understand how genome organization changes during cancer progression, we generated high-resolution (5 kb) genome-wide maps of chromatin contacts using Micro-C in the MCF10A, MCF10AT1, and MCF10CA1a cancer progression series (**Fig. 1A**, **Fig. 1-Fig. Supp. 1A**). We obtained high-quality data with at least 1 billion Micro-C contacts per cell line, spread across 2 biological replicates with 4 technical replicates each (**Supp. File 1**).

**Figure 1.**
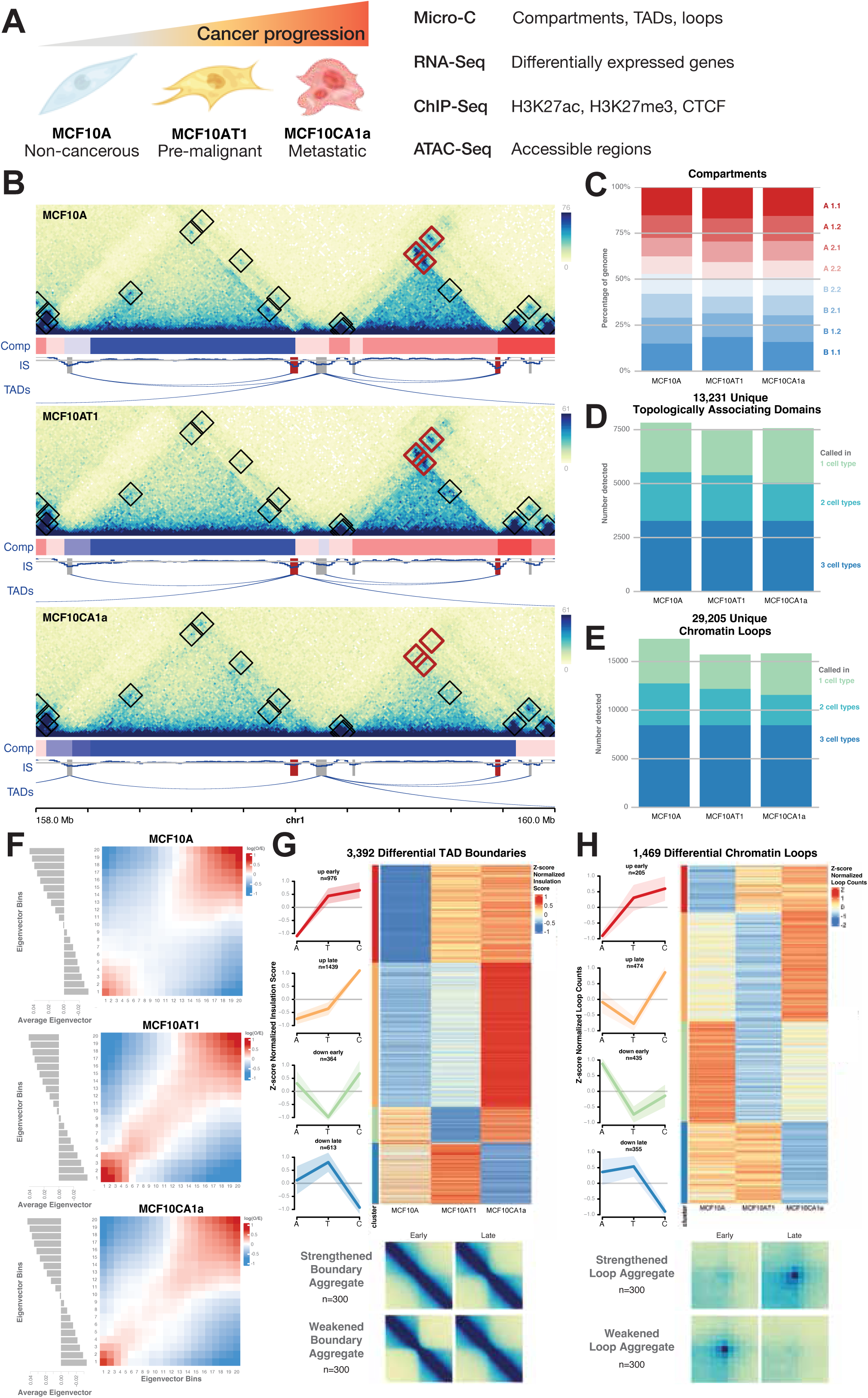
Reorganization of compartments, TADs, and loops during breast cancer progression. **(A)** A diagram of the experimental design. Three epithelial cell lines represent various stages of breast cancer progression; MCF10A are non-cancerous, MCF10AT1 are pre-malignant, and MCF10CA1a are metastatic. In each cell line we generated 5kb resolution Micro-C to identify features such as compartments, topologically associating domains (TADs), and chromatin loops. We overlapped these features with functional changes in gene expression from RNA-Seq, histone modifications and CTCF binding from ChIP-Seq, and chromatin accessibility from ATAC-Seq. **(B)** Micro-C maps of a 2 Mb region of chromosome 1 in MCF10A (non-cancerous), MCF10AT1 (pre-cancerous), and MCF10CA1a (metastatic) cells at 5 kb resolution. Each map has annotations for loop calls, both static (black boxes) and differential (red boxes). Below each map is a track indicating compartment calls from CALDER (dark red is most A-like, dark blue is most B-like) as well as insulation scores tracks with static (grey) and differential (red) boundaries marked. Ribbons indicate TAD calls for each cell type. **(C)** Lengths of the genome assigned to each compartment in each cell type. **(D)** TAD and **(E)** loop calls from each cell type, colored by the number of maps they were initially detected in. **(F)** Saddle plots of interactions between regions of different compartments in MCF10A, MCF10AT1, and MCF10CA1a. Bottom plots indicate the average eigenvector value for each compartment ventile. Plots shown are for chromosomes 2, 12, and 17 (see Methods). **(G)** Left; Differential TAD boundaries clustered by their timing of change, depicted in line plots and heatmap. Right; aggregate plots of weakened and strengthened TAD boundaries (n=100). **(H)** Left; Differential chromatin loops clustered by their timing of change, depicted in line plots and heatmap. Right; aggregate plots of weakened and strengthened loops (n=100).

We identified features of chromatin organization at several levels, including large-scale reorganizations of compartments, medium-scale changes in topologically associating domains (TADs), and fine-scale changes in chromatin loops (**Fig. 1B**). We assigned A and B compartments using CALDER at a resolution of 10kb, TADs using SpectralTAD combined with FAN-C boundary insulation score calculations at 10kb, and loops using SIP at 5- and 10kb resolution (see Methods for details). Each cell type had similar percentages of the genome assigned to the active A (47.1-50.2%) or inactive B (49.7-52.9%) compartment, with the two cancer cell types MCF10AT1 and MCF10CA1a having a slightly higher proportion of A compartment designations (**Fig. 1C**). Similarly, we detected a similar number of TADs (between 7,459-7,825) and chromatin loops (between 15,713-17,332) in each cell line (**Fig. 1D-E**). Although the three cell lines are karyotypically similar due to their shared genetic background, they contain large-scale chromosomal duplications and translocations which were identified by SKY karyotyping analysis (**Fig. 1-Fig. Supp. 1B**), and numerical chromosome aberrations based on SKY and Micro-C sequencing depth analysis were included in the analysis of all chromatin features (see Methods; **Fig. 1-Fig. Supp. 1C, Supp. File 2**). After this correction, chromatin loop counts showed a high degree of reproducibility between technical and biological replicates (**Fig. 1-Fig. Supp. 1D**). We used this deep dataset to characterize structural features that are reshaped during cancer progression.

### Cancer progression reorganizes compartments, TADs, and chromatin loops

Comparative analysis across the three cell types identified significant changes in all major chromatin features during cancer progression (**Fig. 1**, **Fig. 1-Fig. Supp. 2A-B, Supp. File 3-5**).

At a large-scale, we detected changes in compartmentalization. We observed a general shift towards the more active A compartment in early cancer progression, with a larger portion of genomic regions becoming more A-like (31.0%) compared to more B-like (26.0%) in the transition from MCF10A to MCF10AT1, while these changes are more balanced in the later transition from MCF10AT1 to MCF10CA1a (30.0% and 30.6%, respectively). Interactions within the most A-like and B-like compartments were predominant in the pre-cancerous MCF10A cells (**Fig 1F**). However, in MCF10AT1 and MCF10CA1a, stronger interactions appear between more intermediate regions, suggesting a greater degree of intermixing that is consistent with increased compartmental heterogeneity which appears to occur early during cancer progression (**Fig. 1F**) (42,66,67).

TAD boundaries represent genomic regions where upstream and downstream sequences are partially insulated from one another, with fewer contacts between them than within (1,5,19). We detected a total of 13,231 TADs across all three cell types, with 17,097 unique boundaries. TADs detected range in size from 190kb to as large as 3.8 Mb, with a mean of 663kb and a median of 460kb (**Fig. 1-Fig. Supp. 2C**). Assessing changes in insulation score (IS) at TAD boundaries revealed 3,392 (19.8%) boundaries where the degree of insulation changed significantly over the course of cancer progression. Because individual boundaries may be simultaneously used by multiple TADs, the total number of TADs which changed during progression is 5,084 (38.5%) (**Fig. 1G**). There are nearly three times as many boundary changes between later stages in cancer progression (1,693 differential boundaries between MCF10AT1 and MCF10CA1a) than early stages (567 between MCF10A and MCF10AT1). Interestingly, TAD boundaries that gained or lost insulation during progression showed a significant enrichment for weakened boundaries (71.2%) with far fewer boundaries exhibiting increased insulation strength as cancer progressed (28.8%; p-value 0, permutation test, **Fig. 1-Fig. Supp. 2D**). This late-stage weakening of boundaries may reflect a more heterogeneous cell population as cancer progresses (6,68–71).

Chromatin loops are formed by two distal genomic regions that are in more frequent contact than their surrounding or intervening sequences, indicated by higher contact frequency (1,3,10,11). We found 29,205 chromatin loops across all three cell lines, ranging in size from 50 Kb to 2 Mb, with a mean of 402kb and median of 270 Kb in length (**Fig. 1-Fig. Supp. 2C**). 77.6% of loop anchors coincided with CTCF peaks, representing 95.0% of loops with at least one anchor bound by CTCF, and CTCF-bound loops were stronger and longer than non-CTCF loops (**Fig. 1-Fig. Supp. 2E-F**). Loop boundaries often overlapped with TAD boundaries with 52.4% of TADs consisting of loop domains across all cell lines. However, a majority (73.0%) of chromatin loops did not include TADs (**Fig. 1-Fig. Supp. 2G)**. TADs without loop interactions at their boundaries tended to be larger, while loops without TADs can be found at all sizes but are enriched for shorter loops (**Fig. 1-Fig. Supp. 2H**).

To identify loops that changed significantly during cancer progression, we assessed changes in contact frequency among every loop in each cell type, correcting for karyotypic differences that result in differences in coverage between cell lines (see Methods). We identified 1,469 chromatin loops that change significantly over the course of cancer progression, including both weakened and strengthened contacts (**Fig. 1H**), representing 5.0% of all identified loops. Differential loops were defined by a ≥1.5 fold-change between contact frequencies in any two MCF10 cell lines, and an adjusted p-value of <0.025 when considering variation across biological and technical replicates. Unlike TADs, there was a more balanced number of changes between early (1,004 differential loops between MCF10A and MCF10AT1) and late (1,204 between MCF10AT1 and MCF10CA1a) progression stages, as well as between strengthened (679 loops, 46.2% of all differential loops) and weakened loops (790 loops, 53.8% of all differential loops). Interestingly, only a small portion (19.0%) of differential loops were accompanied by changes in CTCF binding (**Fig. 1-Fig. Supp. 2I**). Motif analysis of differential loop anchors revealed only weak motifs of various transcription factors enriched at the boundaries of gained and lost loops, although occupancy did not appear high enough to explain most of the changes we observe (**Fig. 1-Fig. Supp. 2J**). Weakened loops were often associated with a decrease in H3K27ac, a mark of active enhancers, consistent with the notion that active enhancers can help recruit loop extrusion machinery (**Fig. 1-Fig. Supp. 2K**; see below) (72,73).

Taken together, these results demonstrate significant global changes in genome organization during cancer progression across multiple scales from chromatin compartments to loops.

### Comparison to breast cancer patient data

To assess how generalizable the static and dynamic structures detected in the MCF10 progression series are to human tumors, we examined the chromatin loops and TAD boundaries from the MCF10 progression series in non-cancerous mammary epithelial cells (HMEC), five cell lines representing distinct cancer subtypes ranging from less (luminal, HER2+) to more aggressive (triple negative), as well as tissue samples from triple negative breast cancer (TNBC) patients with contralateral normal controls (**Fig. 1-Fig. Supp. 3-4**) (49).

We found that most loops and TAD boundaries detected in MCF10 cells had conserved signatures in each of the other cell types, with chromatin loops generally showing high observed-versus-expected contact frequencies and boundaries showing strong dips in insulation score (**Fig. 1-Fig. Supp. 3A, 4A-B**). When we compared the profiles of differential MCF10 features relative to static MCF10 features within each cell type, we found some cell-specific changes. For example, loops in the “up-early” cluster, which are weaker in MCF10A and stronger in both MCF10AT1 and MCF10CA1a, were significantly stronger than static loops among patient TNBC cells (**Fig. 1-Fig. Supp. 3B**; p-value <= 0.05, Wilcoxon rank sum test). This loop cluster was also stronger in HER2+ (HCC1954) and TNBC A (HCC70) cell lines, and the “down-early” loop cluster is significantly weaker than static loops. Among differential TAD boundaries, those that have strong insulation in MCF10A and MCF10AT1 cells, but reduced insulation in MCF10CA1a, showed stronger insulation among all other examined cancer cell types when compared to static loop boundaries, suggesting that the boundaries that weaken in MCF10CA1a are not necessarily consistently weakened in breast cancer cell lines and tissues (**Fig. 1-Fig. Supp. 4C**; p-value <= 0.05, Wilcoxon rank sum test). These findings highlight that different model systems indeed have distinct profiles of structural change, just as they have distinct gene expression profiles.

### Chromatin loops connect gene promoters to distal regulatory features

We then sought to explore how long-range chromatin interactions relate to gene expression. We identified 17,185 expressed genes across any of the three MCF10 cell types using RNA-Seq and 8,840 differentially expressed genes across all pairwise comparisons in the MCF10 cancer progression (see Methods; **Fig. 2-Fig. Supp. 1A-B, Supp. File 6**). A similar number of genes changed in later stages of cancer development (4,968 between MCF10AT1 and MCF10CA1a) compared to early progression (4,773 between MCF10A and MCF10AT1). Reassuringly, as expected from previous studies (41,64), genes associated with epithelial morphogenesis and cell adhesion were upregulated early during progression, whereas regulation of differentiation, tissue development, metabolism, and signal transduction genes was observed during later stages of progression (**Fig. 2-Fig. Supp. 1C**). These changes are consistent with the development of an intermediate and diverse pre-cancerous state early on during progression, while late changes are known to facilitate metastasis and support the epithelial-to-mesenchymal-like transition observed phenotypically among the progression series (**Fig. 2-Fig. Supp. 1D)** (74–77).

**Figure 2.**
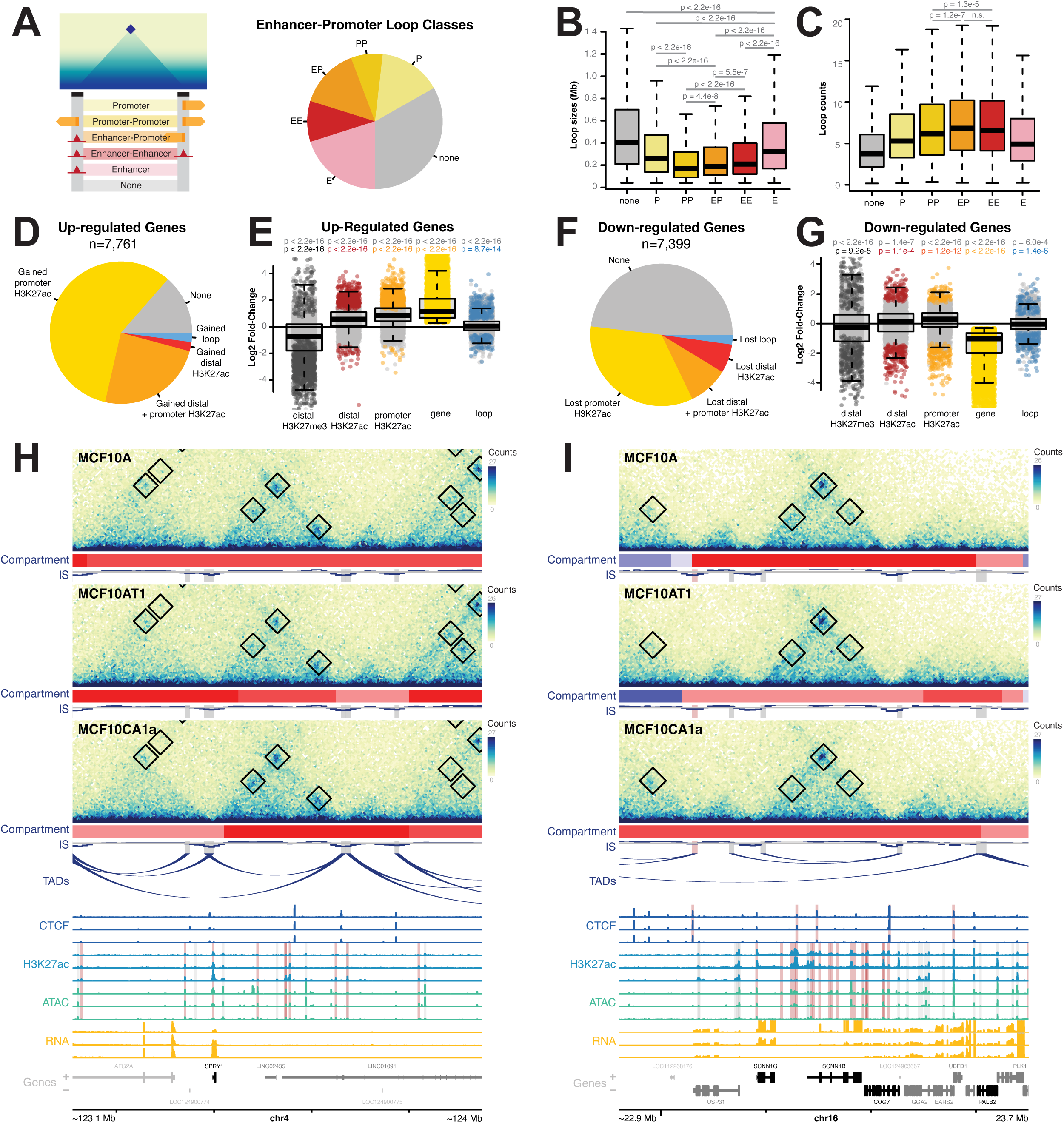
Persistent chromatin loops connect differentially expressed genes to distal enhancers. **(A)** Percentages of loops designated as either promoter-promoter, enhancer-promoter, enhancer-enhancer, or single-sided promoter or enhancer loops. **(B)** Distributions of loop sizes by enhancer/promoter designations. P-values represent Wilcoxon tests comparing the means of different loop classes. Boxplots show the median (middle line), 25^th^ and 75^th^ quartiles (box perimeters), and range excluding outliers (dashed line whiskers). Outliers are defined as values that are over 1.5 times the interquartile range beyond the box bounds and are excluded from these plots. P-values represent Wilcoxon tests comparing the means of various loop sets. Non-significant (n.s.) represents p-values above 0.05. **(C)** Distributions of loop strength by enhancer/promoter designations. P-values represent Wilcoxon tests comparing the means of various loop sets. Non-significant (n.s.) represents p-values above 0.05. **(D)** Percentages of upregulated genes that have gained H3K27ac at promoters, distal enhancers, both, or gained loops. **(E)** Log2 fold-change of distal H3K27me3 (grey), distal H3K27ac (red), promoter H3K27ac (orange), gene expression (yellow), and loop strength (blue), when overlapped. Grey dots indicate features that do not change significantly, while colored points are significantly differential features. Boxplots are defined as in (B). P-values represent Wilcoxon tests comparing the means of each class to 0. Non-significant (n.s.) represents p-values above 0.01. **(F)** Percentages of downregulated genes that have gained H3K27ac at promoters, distal enhancers, both, or gained loops. **(G)** Log2 fold-change of distal H3K27me3 (grey), distal H3K27ac (red), promoter H3K27ac (orange), gene expression (yellow), and loop strength (blue), when overlapped. Boxplot details as defined in (E). **(H)** An example of an upregulated gene (SPRY1) connected to gained enhancers by static loops. Black boxes show loop annotations. Red compartment tracks indicate A compartments, while blue indicates B compartments. In CTCF signal tracks, red highlights indicate differential CTCF peaks. In H3K27ac and ATAC-Seq signal tracks, red highlights indicate differential enhancers as determined by changes in H3K27ac. Genes highlighted in black are differentially expressed. **(I)** An example of downregulated genes (SCNN1G, SCNN1B) connected to lost enhancers by static loops. Plot annotations are as described in (H).

To assess clinical significance of the differentially expressed genes in the MCF10 progression system, we overlapped this gene set with data from the Cancer Genome Atlas breast cancer cohort (TCGA-BRCA) (78,79). A total of 1,884 genes that are differentially expressed between any stages of the MCF10 progression series were also differentially expressed between normal and tumor samples from breast cancer patients, but only roughly half change in the same direction (see Methods; **Fig. 2-Fig. Supp. 1E**). Interestingly, we found a higher degree of directional agreement between late changing genes (i.e. genes that change in MCF10CA1a compared to MCF10A and MCF10AT1) than early changing genes (i.e. genes that change in MCF10AT1 and MCF10CA1a compared to MCF10A). Of note, several looped genes which are differentially expressed in the MCF10 series showed an effect on overall patient survival (**Fig. 2-Fig. Supp. 2A-D)**, and all but one of these genes exhibit significant differences in expression between normal and tumor TCGA-BRCA samples (**Fig. 1-Fig. Supp. 2E**; two-sampled Wilcoxon test of normalized counts, see Methods).

To explore the functional role of chromatin loops, we next overlapped chromatin loops with expressed and differential gene promoters and potential regulatory regions. We identified 52,953 potential enhancers as defined by overlapping histone H3K27ac ChIP-Seq and ATAC-Seq accessibility peaks, commonly used to identify enhancer regions (see Methods) (80–82). Approximately 66.8% of chromatin loops featured some combination of active gene promoters and enhancers within 10kb of loop anchors (**Fig. 2A**). We found that chromatin loops that connect two features (either enhancers or promoters; mean length 260-311 kb) are typically shorter than those that contain only one feature (mean length 376-435 kb) or none (**Fig. 2B**; mean length 532 kb; p-value for all comparisons < 2.2e-16, Wilcoxon rank sum test), with promoter-promoter loops being the smallest on average (mean length 260 kb). Interestingly, enhancer-promoter and enhancer-enhancer loops (mean counts 7.9, 7.9, respectively) were stronger than promoter-promoter loops (mean counts 7.2) despite being longer on average, suggesting that epigenetic signatures associated with active enhancers may support stronger contacts (**Fig. 2C**; p-value for both comparisons < 1.3e-5, Wilcoxon rank sum test).

### Differentially expressed genes show proximal and distal epigenetic changes at persistent chromatin loops

To understand the regulatory modes of action of genes which were differentially expressed during cancer progression, we determined if they had gained or lost the activity-associated H3K27ac mark at their promoters or at distally looped enhancers. We found that while many genes only featured a corresponding change in H3K27ac at their promoter (57.9% of upregulated and 34.1% of downregulated genes), a large percentage also showed changes in distal enhancer activity (26.5% of upregulated and 15.7% of downregulated genes), suggesting that enhancer loops may be playing an important functional role in control of these genes (**Fig. 2D, F**). Comparing the direction of fold-change for genes and promoter H3K27ac, distal H3K27ac, or contact frequency with distal enhancers using Fisher’s Exact test revealed odds ratios significantly higher than 1 for all comparisons (8.8, 2.1, and 1.2, respectively for changes between MCF10A and MCF10CA1a), but that there was a stronger association with promoters than enhancers or loop strength (**Fig. 2-Fig. Supp. 3A-B**). This trend was similar for genes that were differentially regulated both early and late, suggesting that the role of chromatin loops is consistent across all stages of cancer progression. In support of the finding that both distal regulatory changes and changes in contact frequency appear to contribute to changes in gene expression, looped genes that are regulated similarly in MCF10 progression and patient data include both up- and downregulated genes anchored by both static (i.e. *RRM2 and FERMT2)* and dynamic chromatin loops (i.e. *INHBA, PCDH9*; **Fig. 2-Fig. Supp. 3C-F)**.

To further assess the contribution of looped regulatory regions, we explored the degree of change in epigenetic marks and contact frequency between proximal and distal regions. Comparing the changes in acetylation at all gene promoters and distal regulatory regions revealed that upregulated genes exhibit a significant increase in H3K27ac at distally looped enhancers, as well as a significant loss of repressive H3K27me3 marks (**Fig. 2E**; p-value < 2.2e-16, one-sample Wilcoxon test). This trend is weaker for downregulated genes, which feature a closer balance of gained and lost H3K27ac at both enhancers and promoters, as well as both gained and lost H3K27me3 at distal regions (**Fig. 2G**). These results suggest that the static chromatin structures observed during the cancer progression process contribute to the control of differentially regulated genes, particularly among upregulated genes.

We observed that genes that are more strongly expressed are more likely to overlap with the anchor of a chromatin loop, ranging from roughly 33.4% overlap for genes with 0 read counts, 47.0% for genes with 10-1,000 counts, and 56.2% for genes with 5,000 and more counts (**Fig. 2-Fig. Supp. 4A**). To explore whether differential loops are more prominent among genes that change from a low active state to higher expression bins, we analyzed 108 genes that went from an unexpressed or “off” state (2 or fewer read counts) in one cell line to an expressed “on” state (100 or more read counts) in another (**Fig. 2-Fig. Supp. 4B-E**). While these genes were not enriched for differential loops, over 40% overlap with static loops. Similarly, genes that change from a modest “on” state to high expression levels (1000 or more read counts) are not enriched for differential loops, however they do exhibit a higher static loop overlap (61.8%) consistent with higher total gene expression levels (**Fig. 2-Fig. Supp. 4F-I)**. For all gene sets examined, looped genes showed strong and similar trends at distal regulatory regions.

Given that only 5% of loops changed significantly during progression (see Fig. 1), it is not surprising that only a small percentage of differentially expressed genes exhibited significant changes in chromatin contacts with distal enhancers (2.1% of upregulated and 2.2% of downregulated genes; **Fig. 2D, F**). This trend was similar between both early- and late-regulated genes (**Fig. 2-Fig. Supp. 4J**). On average there is a slight but significant change in contact frequency between gene promoters and distal enhancers that corresponds to the change in gene expression (**Fig. 2E-F**). However, most differentially expressed genes are in regions where chromatin structure is essentially stable, reinforcing that persistent structural changes are not universally required for gene regulation.

For example, the *SPRY1* gene, which regulates cell growth and differentiation and has been shown to be upregulated in triple-negative breast cancer tumors (83), is upregulated between MCF10AT1 and MCF10CA1a, and is statically looped to distal enhancers that gain H3K27ac (**Fig. 2H**). Similarly, the *SCNN1G* gene, which encodes for a subunit of a sodium channel and is suppressed in head and neck squamous cell cancer (84), is downregulated between MCF10AT1 and MCF10CA1a, and is statically looped to distal enhancers that lose H3K27ac (**Fig. 2I**). In both cases, the contact frequency remains relatively constant despite changes in distal enhancer acetylation. Many additional examples of statically looped, differentially expressed genes were found, including *RRM2* and *FERMT2* (**Fig. 2-Fig. Supp. 3E-F)**. Taken together, our results show that changes in gene expression are not necessarily accompanied by structural changes, and they suggest that stable chromatin loops may facilitate functionally relevant gene expression programs by providing a pre-existing structure through which differentially regulated distal enhancers can act on target genes.

### Changes in TAD boundary insulation and subcompartments have subtle impacts on gene expression

To explore the impact of changes in insulation at domain boundaries on gene expression, we next examined genes within 50kb of differential boundaries. We found that genes close to weakened boundaries were not enriched for differentially expressed genes, but those near strengthened boundaries were (**Fig. 2-Fig. Supp. 5A**; p-values of 0.141 for early and 0 for late strengthened boundaries, and p-values of 1 for weakened boundaries, permutation test). While strengthened boundaries featured both upregulated and downregulated genes, there was a small but significant correlation between the strength of changing boundaries and fold-change of expression of nearby genes when compared to genes at static boundaries, suggesting an average positive impact on gene expression (**Fig. 2-Fig. Supp. 5B**; p-value 5.9e-3, Wilcoxon rank sum test). This suggests that changes in TAD boundary insulation have small but noticeable impacts on gene expression.

We also examined how subcompartments change genome-wide and at gene promoters. We see that between any two cell types, a majority of subcompartment changes are small changes, i.e. from A.2.2 to A.2.1 (1 step more A-like) or B.1.1 (1 step more B-like), with larger shifts being rarer (**Fig. 2-Fig. Supp. 5C**). The promoters of active genes are enriched for A.1.1 and A.1.2 subcompartments but depleted for all B subcompartments, while inactive gene promoters closely resemble genome-wide distributions (**Fig. 2-Fig. Supp. 5D-E**). The promoters of differentially expressed genes have similar subcompartments at lower and higher expression levels, but these changes are more drastic for genes that shift from on to off or on to high, as defined above (**Fig. 2-Fig. Supp. 5F-H**). Differentially expressed genes with promoters that shift to more B-like by 2 or more subcompartments or more A-like by 3 or more subcompartments have significant impacts on fold-change, but smaller shifts have minimal impacts on gene expression (**Fig. 2-Fig. Supp. 5I**; p-value <= 0.01, Wilcoxon rank sum test). In summary, small changes in subcompartments are very common but appear to have little impact on gene expression, while larger changes correlate more strongly with changes in gene expression during breast cancer progression but are comparatively rare.

### Changes in enhancer acetylation and enhancer-promoter contact are associated with changes in gene expression

To begin to distinguish the effects of enhancer-promoter contact and chromatin looping from enhancer activity effects, we compared gene expression changes at looped and non-looped enhancer-promoter pairs. To do so, we used the activity-by-contact (ABC) model to predict functional enhancer-promoter pairs. ABC combines estimates of enhancer accessibility and activity from ATAC-Seq and H3K27ac ChIP-Seq with enhancer-promoter contact frequency from Micro-C data to generate an estimate of the likelihood of functional enhancer-gene interactions (see Methods) (85). This method allowed us to identify distal regulatory regions that are functionally linked to gene promoters without specifically requiring overlap with chromatin loops. For example, an enhancer and promoter may be in high contact as measured by Micro-C because they overlap with loop anchors, or because they are at close genomic distance. For this analysis, potential enhancers were defined as any overlapping H3K27ac ChIP-Seq and ATAC-Seq peaks 750 bp or farther from the transcription start sites of the target gene.

Applying the ABC model to our data identified 150,056 potential enhancer-promoter pairs across all three cell types. Of these, 53.4% are also promoters of other genes, 23.7% are within the bodies of other genes, and 22.9% are intergenic, and range in distance from 750 to 5M base pairs away from target gene promoters (**Fig. 3-Fig. Supp. 1A**). To better understand the relationship between contact frequency, enhancer activity, and gene expression, we asked how changes in enhancer activity or contact relate to gene expression at target promoters.

**Figure 3.**
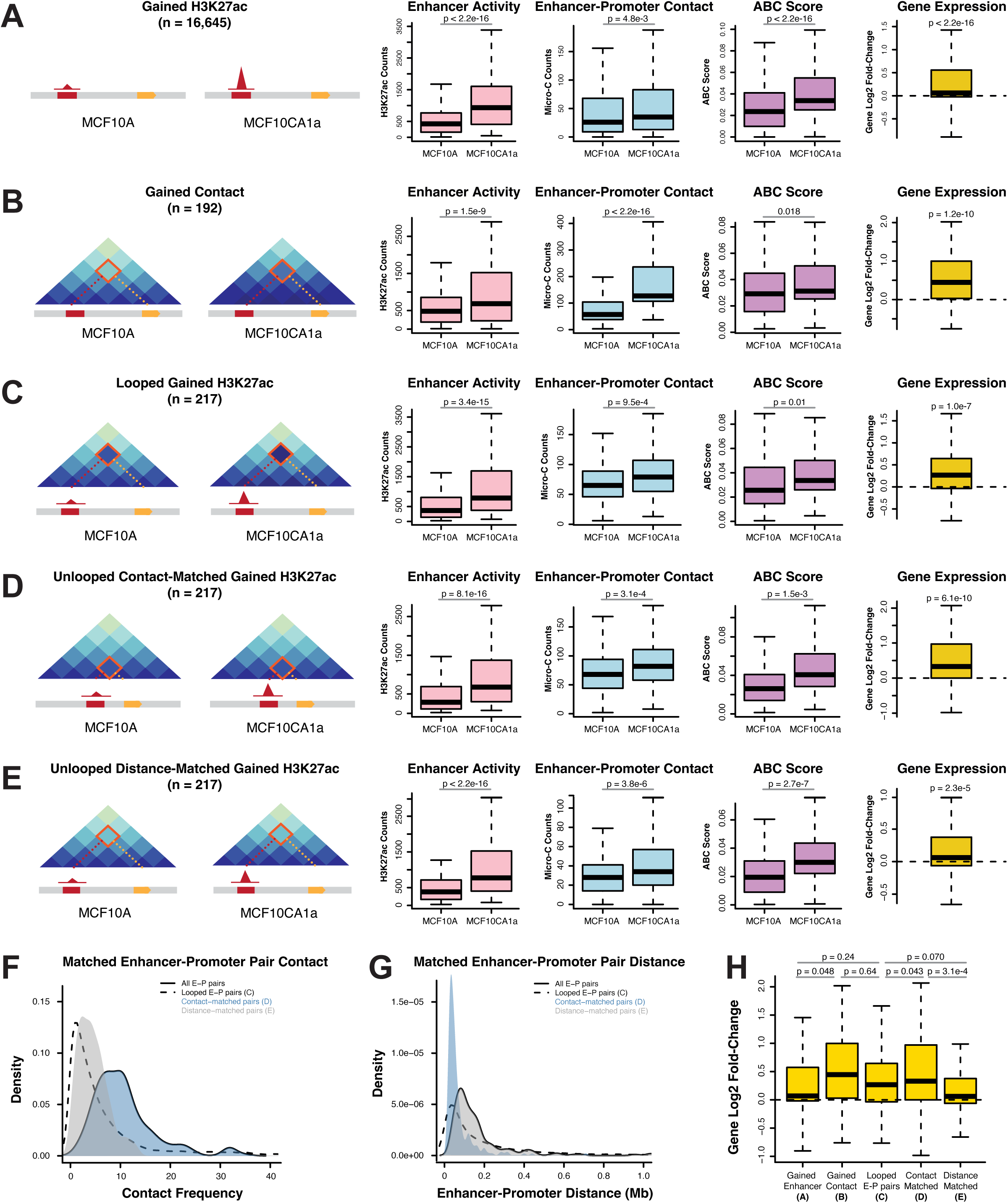
Changes in enhancer acetylation or enhancer-promoter contact are associated with changes in gene expression. Boxplots of distal enhancer H3K27ac (pink), enhancer-promoter contact (blue), and ABC score (purple), as well as gene log2 fold-change (yellow) for enhancer promoter pairs that feature **(A)** differential H3K27ac among enhancers, **(B)** differential enhancer-promoter contact frequency, and **(C)** differential H3K27ac for enhancer-promoter pairs supported by a chromatin loop. Boxplots in **(D)** and **(E)** represent sets of non-looped enhancer-promoter pairs with differential H3K27ac that are matched to the looped set in (C) by contact and distance, respectively. Boxplots show the median (middle line), 25^th^ and 75^th^ quartiles (box perimeters), and range excluding outliers (dashed line whiskers). Outliers are defined as values that are over 1.5 times the interquartile range beyond the box bounds and are excluded from these plots. P-values represent T-tests comparing the means of values in MCF10A and MCF10CA1a for enhancer activity, enhancer-promoter contact, and ABC Score, and T-tests comparing the mean of the value to 0 for gene log2 fold-change. **(F)** Contact distribution of all enhancer-promoter pairs (dashed line), compared to the looped enhancer-promoter pairs in (C, solid line), the contact-matched pairs in (D, blue shade), and the distance-matched pairs in (E, grey shade). **(F)** Distance distribution of all enhancer-promoter pairs (dashed line), compared to the looped enhancer-promoter pairs in (C, solid line), the contact-matched pairs in (D, blue shade), and the distance-matched pairs in (E, grey shade). **(G)** Summary boxplot of the gene log2 fold-change for the enhancer-promoter pairs previously shown in figures (A-E). P-values represent T-tests comparing the means of average gene log2 fold-changes values for different sets of enhancer-promoter pairs.

Pairwise comparison of the MCF10 progression lines indicated that changes in both contact frequency and enhancer activity appear to drive changes in enhancer-promoter networks predicted by ABC **(Fig. 3)**. For example, observing potential enhancers with changes in H3K27ac between MCF10CA1a and MCF10A reveals that these enhancers also exhibit a change in contact frequency and are associated with upregulation of target genes (**Fig. 3A**). We also found that changes in contact frequency are associated with increases in H3K27ac and correlate with higher gene expression (**Fig. 3B**). These results show that not only changes in either contact frequency and enhancer activity correlate with increased gene expression, but they also correlate with each other, suggesting a potentially linked functional role during enhancer-promoter communication.

To then relate enhancer-promoter pairs to chromatin loops and to orthogonally assess whether chromatin loops are acting as a functional bridge for active enhancers, we compared looped and non-looped enhancer-promoter pairs. Enhancer-promoter pairs that have changes in distal H3K27ac and are supported by chromatin loops correlated with changes in gene regulation (**Fig. 3C**). This effect was stronger than distance-matched non-looped enhancer-promoter pairs, but similar to contact-matched non-looped pairs, suggesting that increased contact frequency caused by loop extrusion may contribute to the stronger correlation with gene expression (**Fig. 3D-E**). Contact-matched non-looped pairs were closer on average to the looped pairs of similar contact frequency, while distance-matched non-looped pairs were in less-frequent contact than looped pairs of similar genomic distance (**Fig. 3F-G**).

Comparing the distributions of target gene fold-change for these various sets of enhancer-promoter pairs revealed several trends (**Fig. 3H**). First, pairs with significant changes in contact have a larger mean gene fold-change than pairs with significant changes in activity, suggesting that either can contribute to changes in enhancer-promoter functional pairing but that contact may have a particularly strong impact. Second, looped enhancer-promoter pairs have a comparable or larger mean gene fold-change to pairs with changes in activity or contact, suggesting again that chromatin loops may support functional enhancer-promoter pairs. Lastly, looped pairs have a similar mean gene fold-change as contact-matched pairs, which in turn have a higher mean gene fold-change than distance-matched pairs, suggesting that the increased contact frequency that chromatin loops provide to enhancer-promoter pairs may be a driving force for the functional connection. These trends hold true for all tested pairwise comparisons between cell types (**Fig. 3-Fig. Supp. 1B-C**).

Taken together, these findings demonstrate that not all gene regulatory changes are accompanied by chromatin reorganization, but when it occurs reorganization appears to facilitate functional changes.

### Changes in chromatin looping are enriched for progression-associated differentially expressed genes

We next explored how changes in chromatin looping may functionally contribute to gene regulation during cancer progression. To do so, we compared changes in chromatin loop contacts to alterations in expression of progression-associated genes. Overall, while only a small subset of gene promoters overlaps with the anchors of differential chromatin loops (507 genes, 3.0% of expressed genes), those that do are significantly enriched for genes that are differentially expressed during cancer progression based on permutation analysis (331 differentially expressed genes, 65.3% of all differentially looped genes; **Fig. 4-Fig. Supp. 1A**).

**Figure 4.**
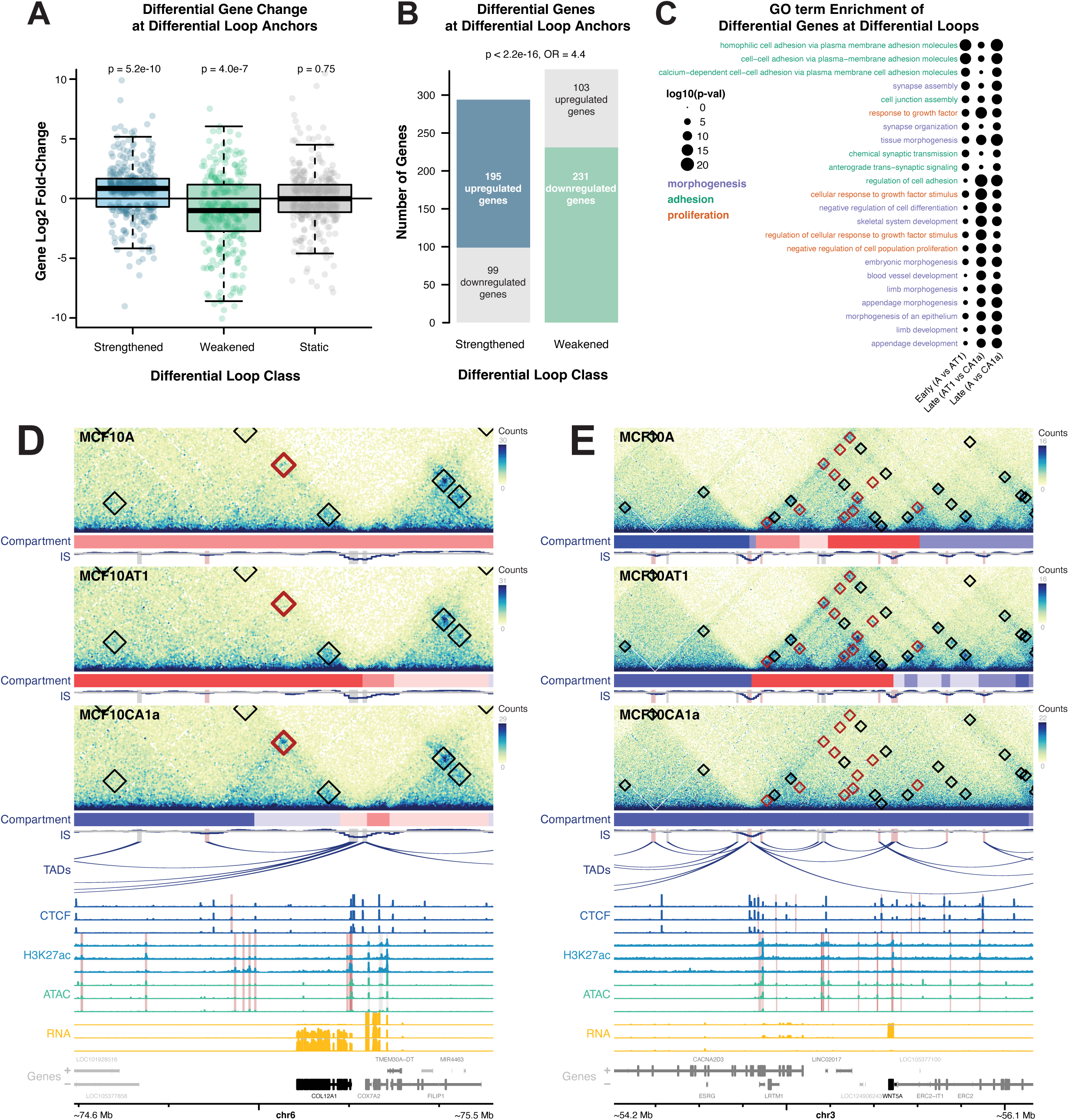
Differential loops are enriched for cancer-relevant differentially expressed genes. **(A)** Log2 fold-change of differentially expressed genes at the anchors of gained (blue), weakened (green), or static (grey) loops. Boxplots show the median (middle line), 25^th^ and 75^th^ quartiles (box perimeters), and range excluding outliers (dashed line whiskers). Outliers are defined as values that are over 1.5 times the interquartile range beyond the box bounds and are excluded from these plots. P-values represent Wilcoxon tests comparing the mean of each set to 0. **(B)** Bar plot showing the number of differentially expressed genes at strengthened or weakened loop anchors. Bar segments are colored by whether the gene is changing the same (blue for upregulated in strengthened loops, green for downregulated in weakened loops) or opposite (grey) direction as the loop. P-value represents a Fisher’s Exact Test for whether the odds ratio (OR) is greater than 1. **(C)** GO term enrichments for genes upregulated in MCF10A, MCF10AT1, or MCF10CA1a. Size indicates p-value. Terms are color-coded based on gene type; morphogenesis (purple), proliferation (orange), and cell adhesion (teal). **(D)** An example of an upregulated gene (COL12A1) with a promoter that overlaps a strengthened loop with distal enhancers. Black boxes show loop annotations, while red boxes indicate differential loops. Red compartment tracks indicate A compartments, while blue indicates B compartments. In CTCF signal tracks, red highlights indicate differential CTCF peaks. In H3K27ac and ATAC-Seq signal tracks, red highlights indicate differential enhancers as determined by changes in H3K27ac. Genes highlighted in black are differentially expressed. **(E)** An example of a downregulated gene (WNT5A) with a promoter that overlaps with several weakened loops containing distal enhancers that lose H3K27ac. Plots are annotated as in (A).

We asked whether there was a relationship between the formation or loss of loops and differential expression of these loop-associated genes. We indeed found that differential loops are more likely to change in the same direction as the gene (i.e. increased contact frequency with distal regions associated with increased gene expression) (**Fig. 4A-B**). The fold-change of differential expressed genes which also showed differential loping were significantly higher than a random sampling of differentially expressed genes (**Fig. 4A**). For example, of 3,261 genes that were differentially upregulated between MCF10A and MCF10CA1a, loops were significantly strengthened at 98 of these genes and significantly weakened at 31. Similarly, of 3,088 downregulated genes, 65 genes overlap weakened loop anchors and 41 genes overlap strengthened loop anchors. In contrast, the number of expected genes at strengthened or weakened loops for a random sampling of genes this size is 38 and 32, respectively (**Fig. 4-Fig. Supp. 1B-C**). We also found a subset of chromatin loops and genes changed in opposite directions (**Fig. 4B**). The genes whose changes in expression correlate with changes in looping are enriched for several cancer-relevant pathways, such as morphogenesis, differentiation, and proliferation (**Fig. 4C**) (86).

In total, we identified 127 unique genes upregulated in areas that experience increased chromatin contacts, either at loop anchors or within existing structures. As an example, the promoter of the *COL12A1* gene, which encodes a collagen protein known to be associated with aggressive and mesenchymal phenotypes, overlaps a loop boundary that is very weak in MCF10A cells where *COL12A1* is not expressed (87–89). As *COL12A1* gene expression is upregulated during progression, contacts increase at a distal region 310 kb away, and H3K27ac and accessibility also increase at these likely distal regulatory regions **(Fig. 4D).**

Similarly, we observe 123 unique genes that are downregulated during oncogenic progression. One example is *WNT5A* which encodes for an important signaling protein and is downregulated in breast cancer and across MCF10A progression (90–92). Similar to *COL12A1*, the promoter of *WNT5A* is involved in many differential distal regulatory contacts, ranging in distance from 240 to 640 kb (**Fig. 4E**). Unlike *COL12A1*, these contacts are strongest in MCF10A cells and severely weaken in MCF10AT1 and MCF10CA1a cells. Accompanying these changes in contact, the distal regulatory regions that appear to support *WNT5A* in MCF10A cells become deacetylated and decrease in accessibility. Another example of an upregulated genes that is looped to a distal enhancer is *RASA1*, which gains contact frequency as gene expression increases, while *ZEB2* and *SDC3* are downregulated genes that lose enhancer contact frequency similar to *WNT5A* (**Fig. 4-Fig. Supp. 2A-C**). Other loci showed the opposite behavior; for example, *TNFRSF21* and *HS3ST3A1* are upregulated genes at the anchors of weakened loops, while *NNMT* is a downregulated gene at a weakened loop anchor (**Fig. 4-Fig. Supp. 2D-F**).

### Genome reorganization occurs at cancer-relevant genes and is accompanied by proximal and distal changes in histone marks

Finally, we aimed to comprehensively explore the genes that are differentially regulated in areas that also have strong changes in chromatin looping, to better understand the potential regulatory mechanisms at play.

A locus-by-locus view of gene and loop fold-change allows us to view the relationship between changes in expression and structure among each pairwise comparison of cells (**Fig. 5A-C**). While we see that a majority of genes have a corresponding change in looping (i.e. up-regulated genes overlapping strengthened loops), we observed that the percentage of corresponding changes increases in the later stages of cancer progression. For example, the percentage of differential loop-gene pairs where the gene overlaps at least one gained loop is 47.5% and 69.9% among genes up-regulated in MCF10A compared to MCF10AT1 and MCF10CA1a, respectively, 66.7% and 79.6% among genes up-regulated in MCF10AT1 compared to MCF10A and MCF10AT1, respectively, and 70.7% and 58.0% among genes up-regulated in MCF10CA1a compared to MCF10A and MCF10AT1, respectively. This may indicate that the regulatory impacts of changes in chromatin looping occur over longer timescales, or that genes impacted by changes in chromatin structure may be more common in metastatic cells. We did not, however, find any correlation between the magnitude of loop fold-change and gene fold-change.

**Figure 5.**
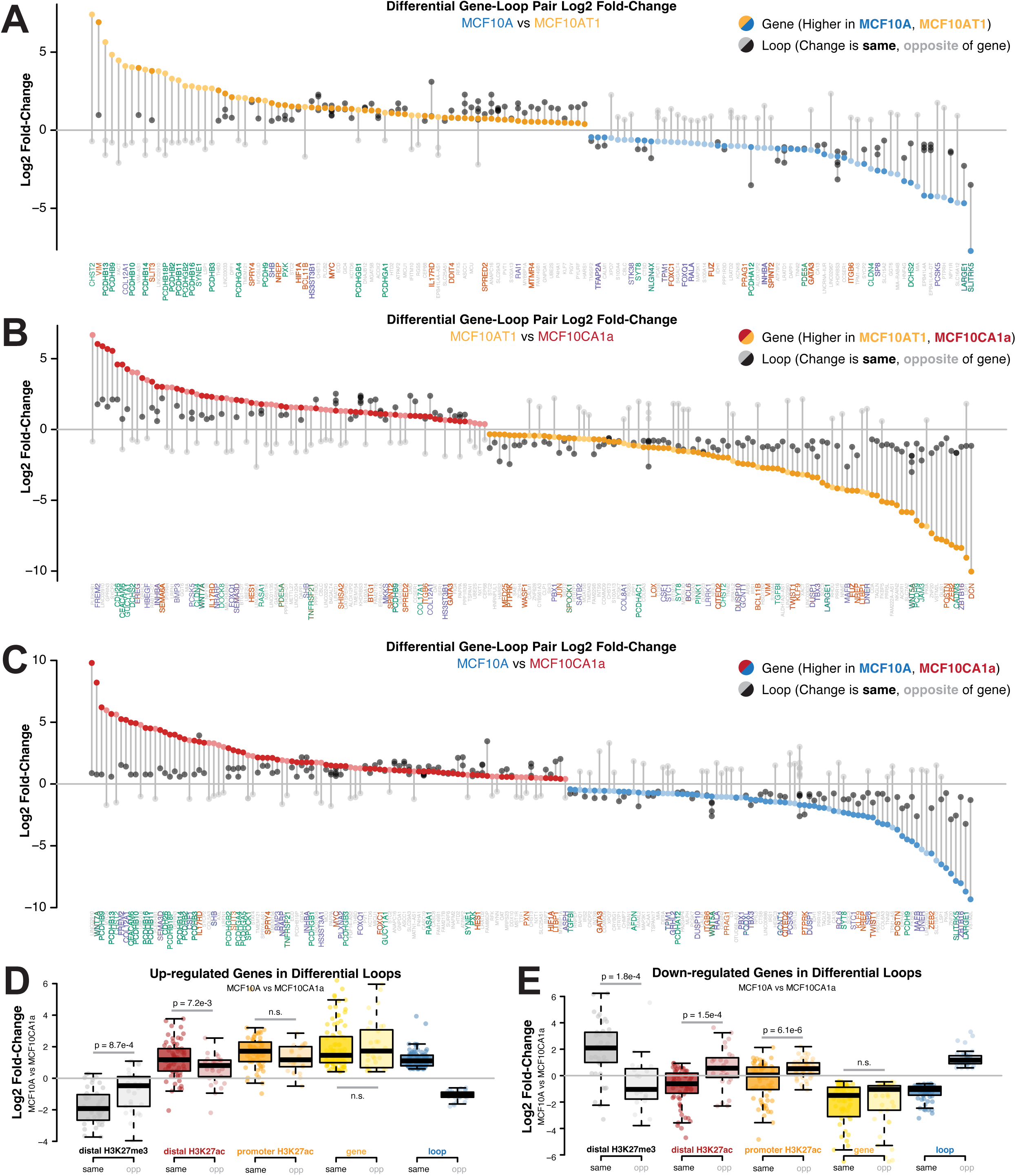
Progression-associated differentially expressed genes exhibit local and distal epigenetic changes at differential loops. (A-C) Log2 fold-change of genes (colored dots) and the differential loops they overlap with (black/grey dots) for genes and loops that change between **(A)** MCF10A and MCF10AT1, **(B)** MCF10AT1 and MCF10CA1a, and **(C)** MCF10A and MCF10CA1a. Gene labels are below. **(D)** Log2 fold-change between MCF10A and MCF10CA1a of distal H3K27me3 (grey), distal H3K27ac (red), promoter H3K27ac (orange), gene expression (yellow), and loops (blue) among upregulated genes that overlap with gained loops (darker colors) or lost loops (lighter colors). Boxplots are defined as in (A). P-values represent T-tests comparing the mean values of the features at loops that change in the same and those that change in opposite directions from the differential genes at their anchors. Non-significant (n.s.) p-values are any p- values above 0.01. **(E)** Log2 fold-change of distal H3K27me3 (grey), distal H3K27ac (red), promoter H3K27ac (orange), gene expression (yellow), and loops (blue) among downregulated genes that overlap with gained loops (darker colors) or lost loops (lighter colors). P-values represent T-tests comparing the mean values of the features at loops that change in the same and those that change in opposite directions from the differential genes at their anchors. Non-significant (n.s.) p-values are any p-values above 0.01.

To better understand how changes in looping are related to gene expression, we compared patterns of gene expression, promoter H3K27 acetylation, and distal enhancer H3K27 acetylation or trimethylation at looped genes that change in either the same or opposite direction as the loops they overlap (**Fig. 5D-E**, **Fig. 4-Fig. Supp. 1D-E**). These two histone H3 modifications are mutually exclusive and have opposite effects on gene expression, marking enhancers and silencers, respectively (93). Both modifications are able to impact distal gene expression via chromatin interactions (94).

Up-regulated genes between MCF10A and MCF10CA1a showed similar epigenetic signatures at both proximal and distal regions, regardless of whether the loop they overlap gets stronger or weaker (**Fig. 5D**). Promoter regions gained H3K27ac, while distal regions gained H3K27ac and lost H3K27me3. There is however a significant difference in the extent of distal changes depending on the loop direction, with strengthened loops exhibiting a significantly higher increase in distal H3K27ac and a decrease in H3K37me3 marks. This behavior further supports the notion that these distally looped regulatory regions are important functional elements. Down-regulated genes showed distinctly different signatures (**Fig. 5E**). Genes that overlap with weakened loop anchors showed decreased promoter H3K27ac and distal H3K27ac and increased distal H3K27me3, consistent with signatures typically associated with reduced gene expression (93). Interestingly, genes that overlap strengthened loop anchors showed different patterns, with a gain in promoter H3K27ac and loss of distal H3K27me3 repressive marks. We conclude that expression of a subset of progression-associated genes is strongly correlated with loop formation. These trends are similar but weaker for genes that change between different cell types (**Fig. 4-Fig. Supp. 1D-E**).

Taken together, our genome-wide analysis of structural and regulatory changes during MCF10A cancer progression has highlighted hundreds of restructured regions where cancer-relevant genes are differentially regulated. These findings suggest that, while relatively rare, changes in chromatin looping may be associated with regulatory changes that drive expression of hundreds of oncogenes during cancer progression.

## DISCUSSION

Multiple levels of chromatin organization and integration with epigenetic parameters contribute to regulation of gene expression. We have identified dynamic chromatin organizational changes on multiple scales across breast cancer progression using the MCF10A model system. By comparing both a pre-malignant and metastatic cell line to non-cancerous epithelial cells we were able to detect both early- and late-stage changes as well as similarities in genome structure and relate them to gene expression changes, including many cancer-relevant genes.

We found that compartmental shifts occur more often in early stages of cancer development. This behavior is consistent with previous studies that have shown intermixing of chromatin A and B compartments in cancer cells (42,50). The general shift towards more active compartments during cancer progression may reflect a broader epigenetic landscape and greater heterogeneity in gene expression within cancer cell populations (6,67,95–97).

On a finer scale, structural changes in TADs occurred more often during later stages of metastasis. We also found an abundance of weakened TAD boundaries in MCF10 breast cancer progression compared with boundaries that gain insulation. This finding is in line with previous studies which have shown TAD weakening in triple-negative breast cancer (49,70), but is contrary to other observations which have suggested that prostate cancer is associated with the formation of many additional TADs (46), or findings of minimal changes in TAD structure in colorectal or breast cancer (42,50). The differences in TAD changes across cancer studies may reflect differences in the types of cancer samples analyzed or the analysis methods used. For example, we detected subtle changes in insulation score at TAD boundaries in the MCF10 model, while other methods tailored towards detecting more drastic restructuring events such as the formation of TAD cliques or TAD fusions found different behaviors (50,71). Recent studies comparing chromatin structures across patient samples have also revealed a high degree of TAD and loop heterogeneity among tumors even of the same cancer type, and it is thus not surprising that patterns observed between different cancer types and in a cell culture model could vary even further (71). The weakened TAD boundaries in the MCF10 progression system are typically strong boundaries in cancer types, suggesting that the same boundaries are not weakened universally in cancer. The weakening of TAD boundaries is intriguing in that it may point to higher heterogeneity in chromatin structure in more aggressive cancer stages, possibly contributing to more extensive gene misregulation (67,70,71). Further studies using single-cell analysis in the MCF10CA1a population will be necessary to confirm if this is the case.

Comparing the structural and functional features of the MCF10 progression system to other breast cancer datasets revealed many common chromatin loops, TADs, and differentially expressed genes, as well as differences between various cancer progression models and patient data. There are hundreds of genes that change over the course of MCF10 progression that are also significantly different in breast cancer tumors from patients compared to healthy controls, including many that overlap with chromatin loops; however, this represents only roughly a quarter of all differential genes in the MCF10 progression model. Despite this transcriptional diversity, chromatin loops and TAD boundaries detected in MCF10 cell lines showed similar average signatures in mammary epithelial cell lines and triple-negative breast cancer patients. However, cell type-specific differences do appear among differential features based on their timing and direction of change in the MCF10 progression model. For example, loops that are stronger in metastatic MCF10CA1a cells were also significantly stronger than static MCF10 loops among TNBC patients, making them a particularly interesting subset of regions to explore further.

Chromatin loops functionally connect gene promoters to distal regulatory elements. In the MCF10 model, many genes differentially regulated during cancer progression are associated with chromatin loops shared between all three cell lines, but which show changes in distal enhancer H3K27ac and H3K27me3. These trends were stronger with up-regulated genes, suggesting that we may need to explore different epigenetic signatures to better understand how chromatin structure may influence repression. We also observed that more highly expressed genes are more likely to overlap with chromatin loops. Activity-by-contact analysis showed that looped enhancer-promoter pairs also exhibit greater correlation between distal enhancer H3K27ac and gene expression than non-looped enhancers due to their increased contact frequency. These findings suggest that persistent chromatin loops that do not change during cancer progression nevertheless have functional relevance and that they do so by bridging enhancers to target gene promoters.

Activity-by-contact analysis also revealed that subtle changes in chromatin contact can contribute to the rewiring of enhancer-promoter regulatory connections. Enhancer-promoter pairs that exhibit changes in contact correlated with stronger changes in target gene expression than those with only changes in activity, in line with the concept that contact with distal regulatory elements is an important component of gene regulation (98). We also found that changes in chromatin contact are associated with more modest changes in activity, and vice-versa. This correlation between enhancer activity and enhancer-promoter contact further points to a functional link between the two. Together, these results suggest that both contact frequency and activity contribute to enhancer-promoter connectivity.

Strong changes in contact at chromatin loops are relatively rare across cancer progression, but many clear examples do exist, and they are notably enriched for differentially expressed genes. Interestingly, only a small portion of differential loops can be explained by changes in CTCF binding at anchors, suggesting other forces may be influencing their contact frequency. Loss of H3K27ac can be seen at the anchors of lost loops, consistent with the idea that active enhancers can help recruit cohesin to chromatin (72,73). Therefore, some weakened loops might be explained by a loss of H3K27ac which leads to a loss of cohesin at that region.

We also found that genes at differential loops are more likely to be differentially regulated in the same direction as the change in loop strength. This supports the notion that a subset of genes may be regulated in a structure-dependent manner (71,99,100). This interpretation is in line with previous observations which have shown a subset of genes to be sensitive to cohesin or NIPBL depletion which disrupts chromatin loops (11,17,20,25,32,101–103). Importantly, these findings suggest this subset of structure-sensitive genes may include many with relevance to breast cancer progression.

Epigenetic signatures at gene promoters and distal regions differed based on the direction of gene change. Up-regulated genes consistently showed a gain of active-associated H3K27ac marks at both promoters and distal regions, and a loss of distal repressive H3K27me3 marks. These changes were shared between static, strengthened, and weakened loops, although up-regulated genes at strengthened loops had stronger distal changes. These findings are consistent with the loops functionally supporting interactions with distal enhancers via increase contact. However, down-regulated genes showed more complex patterns. Fewer down-regulated genes could be explained by changes in H3K27ac or H3K27me3 at the promoter or distal regions, with over half of down-regulated genes having no clear epigenetic driver, compared to only 15% of up-regulated genes. This suggests that these histone marks are not sufficient to explain down-regulated genes as well as they can explain up-regulated genes. Down-regulated genes at differential loops also showed opposite patterns based on the direction of loop change; weakened loops showed loss of distal H3K27ac and gain of H3K27me3 consistent with an inactivated enhancer, while strengthened loops showed the opposite. Additional studies will be required to fully understand the repressive mechanisms in this system and how they relate to chromatin structure.

Our study has several limitations. While the MCF10 progression system is a well-established and -characterized model, it does not reflect the complexities of in vivo tumor formation. In addition, the MCF10 cell lines used here do not fully represent all stages of cancer progression as it encompasses normal, premalignant and metastatic states, but lacks a separate malignant but non-metastatic state such as MCF10DCIS. Furthermore, based on the largely correlative nature of our analysis, we are unable to determine the causal relationships of chromatin structure changes at the loci of differentially expressed genes. Follow-up studies exploring any of the gene loci identified here as possibly being contact-dependent could help elucidate these relationships. Additionally, as a population-level assay, Micro-C is only able to provide aggregate data across an entire population of cells. To address how cell-to-cell heterogeneity contributes to some of the functional relationships we observe, and whether that heterogeneity is occurring at a cellular or population level, we would need to apply single-cell sequencing or imaging-based approaches. These questions will be the subjects of future studies.

In summary, using the MCF10 breast cancer model, we have generated a rich genomic dataset of structural and functional changes in the genome during breast cancer progression. Our data uncovers new insights into the structure function relationship in gene regulation and into the role of genome organization during malignant breast cancer progression.

## MATERIALS AND METHODS

| Key Resources Table |  |  |  |  |
| --- | --- | --- | --- | --- |
| Reagent type (species) or resource | Designation | Source or reference | Identifiers | Additional information |
| cell line (*Homo sapiens*) | Cell line MCF10A | ATCC | Cat#:CRL-10317; RRID:CVCL_0598 | Non-cancerous mammary epithelial cells |
| cell line (*Homo sapiens*) | Cell line MCF10AT1 | Karmanos Cancer Institute | ID:MCF10AneoT; RRID:CVCL_5554 | Pre-malignant mammary epithelial cells |
| cell line (*Homo sapiens*) | Cell line MCF10CA1a | Karmanos Cancer Institute | ID:MCF10CA1a.cl1; RRID:CVCL_6676 | Metastatic mammary epithelial cells |
| antibody | anti-human CTCF (rabbit monoclonal) | Cell Signaling Technology | Cat#:3418 | ChIP-Seq (20 ul per 150 ug chromatin) |
| antibody | anti-human H3K27ac (rabbit polyclonal) | AbCam | Cat#:ab4729; RRID:AB_2118291 | ChIP-Seq (2 ug per 10 ug chromatin) |
| antibody | anti-human H3K27me3 (mouse | AbCam | Cat#:ab6002; RRID:AB_305237 | ChIP-Seq (4 ug per 15 |
|  | monoclonal) |  |  | ug chromati<br>n) |
| commerci<br>al assay<br>or kit | Dovetail<br>Micro-C Kit &<br>Assay Service | Dovetail Genomics | Cat#:20101E | Loop<br>level<br>project |
| commerci<br>al assay<br>or kit | Direct-zol<br>RNA Miniprep | Zymo Research | Cat#:R2050 |  |
| commerci<br>al assay<br>or kit | NEBNext rRN<br>A Depletion<br>Kit v2 | New England Biolabs | Cat#:7400 |  |
| commerci<br>al assay<br>or kit | NEBNext<br>Ultra II RNA-Seq Library<br>Prep Kit | New England Biolabs | Cat#:E7770 |  |
| software,<br>algorithm | BWA mem | PMID: 19451168 | RRID:SCR_0109<br>10 | Version<br>0.7.17 |
| software,<br>algorithm | Pairtools | DOI:<br>10.1101/2023.02.13.528<br>389 | RRID:SCR_0230<br>38 | Version<br>0.3.0 |
| software,<br>algorithm | samtools | PMID: 33590861 | RRID:SCR_0021<br>05 | Version<br>1.17 |
| software,<br>algorithm | Juicer tools | PMID: 27467249 | RRID:SCR_0172<br>26 | Version<br>1.22.01 |
| software,<br>algorithm | NeoLoopFind<br>er | PMID: 34092790 |  |  |
| software,<br>algorithm | CALDER | PMID: 33972523 |  | Version<br>2.0 |
| software,<br>algorithm | FAN-C | PMID: 33334380 |  | Version<br>0.9.21 |
| software,<br>algorithm | SpectralTAD | PMID: 32689928 |  | Version<br>1.18.0 |
| software,<br>algorithm | SIP | PMID: 32127418 |  | Version<br>1.6.2 |
| software,<br>algorithm | STAR | PMID: 23104886 | RRID:SCR_0044<br>63 | Version<br>2.4 |
| software,<br>algorithm | Trim Galore | DOI:<br>10.5281/zenodo.512789<br>8 | RRID:SCR_0118<br>47 |  |
| software,<br>algorithm | Bowtie2 | PMID: 22388286 | RRID:SCR_0163<br>68 |  |
| software,<br>algorithm | MACS2 | PMID: 18798982 | RRID:SCR_0132<br>91 |  |
| software,<br>algorithm | DESeq2 | PMID: 25516281 | RRID:SCR_0156<br>87 | Version<br>1.42.0 |
| software,<br>algorithm | CSAW | PMID: 26578583 | RRID:SCR_0267<br>38 |  |
| software,<br>algorithm | GEPIA | PMID: 40396370 | RRID:SCR_0282<br>42 |  |
| software,<br>algorithm | GenomicRanges | PMID: 23950696 | RRID:SCR_0000<br>25 |  |
| software,<br>algorithm | mariner | PMID: 38814811 |  |  |
| software,<br>algorithm | plotgardener | PMID: 35134826 |  |  |
| software, algorithm | Activity-by-contact | PMID: 31784727 |  |  |
| software, algorithm | nullranges | PMID: 37084270 |  |  |
| software, algorithm | GSEA | PMID: 16199517 | RRID:SCR_003199 |  |
| software, algorithm | HOMER | PMID: 20513432 | RRID:SCR_010881 |  |

### Cell culture

MCF10A cells were obtained from AATC (CRL-10317). MCF10AT1 (MCF-10AneoT) and MCF10CA1a (MCF10CA1a.cl1) cells were obtained from the Karmanos Cancer Institute. All cells were received frozen, thawed in media and grown to a confluence of 80% after 2-5 passages. Stock solutions were frozen down to be used for all experiments. The identity of each cell line has been authenticated using short tandem repeat profiling and tested periodically for the absence of mycoplasma.

MCF10A and MCF10AT1 cells were cultured in standard high calcium medium with growth factors, consisting of DMEM/F12 (Invitrogen, 21041025) supplemented with 1.05 mM CaCl2, 10 mM HEPES, 10 ug/ml insulin (Sigma, I1882), 20 ng/ml EGF (Peprotech, AF-100-15), 0.5 ug/ml Hydrocortisone (Sigma, H0135), 100 ng/ml cholera toxin (Sigma, C8052), and 5% horse serum (Invitrogen, 16050122). MCF10CA1a cells were cultured in standard high calcium medium without growth factors, consisting of DMEM/F12, 1.05 mM CaCl2, 10 mM HEPES, and 5% horse serum.

### Karyotypic analysis

Metaphase chromosomes were prepared by incubating cells with 100 ng/ml Colcemid (Roche, Brighton, MA) for two hours, followed by mitotic shake-off. The mitotic cells were then treated with a hypotonic solution (0.075M KCl) for 20 minutes at 37°C. After this treatment, the cells were centrifuged, the supernatant was extracted, and the cells were initially fixed with a methanol/acetic acid solution (3:1). This step was repeated three times. Finally, the cells were fixed onto a slide using a humidity-controlled Thermotron (Thermotron, Holland MI).

SKY probes were purchased from Applied Spectral Imaging (Carlsbad, CA) and hybridized onto slides that were aged at 37°C for 3 days. Detection was then carried out according to the protocol provided by Applied Spectral Imaging, using the following purchased antibodies: Avidin Cy5 (Rockland Immunochemicals, Limerick, PA), Mouse Anti-Digoxin antibody (Sigma-Aldrich), and a goat anti-mouse antibody conjugated to Cy5.5 (Rockland Immunochemicals). Slides were then mounted and counterstained with DAPI (Vector Laboratories, Newark, CA). Hybridization occurred over a period of two days at 37°C. Approximately 20-25 metaphases were imaged and karyotyped using the ASI GenASIS 8.2.2 software on an Olympus BX63 microscope (Evident, Tokyo, Japan) equipped with a Spectral Cube (Applied Spectral Imaging, Carlsbad, CA).

Composite karyotypes for the three cell lines, listing abnormalities found in three or more metaphases, are as follows:

**MCF10A**: 41-53, XX, idic(1)(q10), +1, der(3)t(3;9)(p21;p22), i(8)(q10), der(9)t(3;5;9)(p14;p23;q34;p21)

**MCF10AT1**: 42-49, XX, der(3)t(3;9)(p21;p22), t(3;17)(p14;p11.2), t(6;19)(p25;q12q13.3), der(9)t(3;5;9)(p14;p23;q34;p21), t(19;9)(q34;p11),+20

**MCF10CA1a**:43-63, XX, der(3)t(3;9)(p21;p22), +der(3)t(3;9)(p21;p22), t(3;17)(p14;p11.2), der(9)t(3;5;9)(p14;p23;q34;p21), +10

### Micro-C library preparation

Cells were frozen and pellets of 1M cells were used for Micro-C library preparation. The Micro-C library was prepared using the Dovetail® Micro-C Kit according to the manufacturer’s protocol. Briefly, the chromatin was fixed with 0.3M disuccinimidyl glutarate (DSG) and 37% formaldehyde in the nucleus. The cross-linked chromatin was then digested in situ with micrococcal nuclease (MNase). Following digestion, cells were lysed with 20% SDS to extract the chromatin fragments and the chromatin fragments were bound to Chromatin Capture Beads. Next, the chromatin ends were repaired and ligated to a biotinylated bridge adapter followed by proximity ligation of adapter-containing ends. After proximity ligation, the crosslinks were reversed, the associated proteins were degraded, and the DNA was purified then converted into a sequencing library using Illumina-compatible adaptors. Biotin-containing fragments were isolated using streptavidin beads prior to PCR amplification. Each library was sequenced on an Illumina Novaseq platform to generate 240-340 million 2 x 150 bp read pairs (average depth of 280M reads). Mico-C libraries were prepared by Dovetail Genomics (Scotts Valley, CA). Four pellets and subsequent Micro-C libraries (technical replicates) were prepared per culture of cells (biological replicates), with two biological replicates grown per cell line on separate weeks.

### RNA-Seq library preparation

Total RNA was isolated from cells using Trizol (Life Technologies) and purified using the Direct-zol RNA kit (Zymo Research, Irvine, CA, USA: R2050). RNA quality and quantity were assessed using the RNA 6000 Nano Kit with the Agilent 2100 Bioanalyzer (Agilent Technologies, Santa Clara, CA). RNA quantity was further assessed using a Nanodrop2000 (Thermo Scientific, Lafayette, CO) and Qubit HS RNA assay (Thermo Fisher Scientific). Total RNA was depleted of ribosomal RNA (New England Biolabs, NEB #7400), reverse transcribed using a random hexamer strategy, and strand-specific adapters were added following the NEBNext Ultra II RNA-seq library prep kit (New England Biolabs, E7770). Paired-End sequencing was used to generate high depth coverage ranging from 80 to 120 million paired-end reads. Three biological replicates were generated per cell line from independent cultures and library preparations.

### ChIP-Seq library preparation

ChIPseq for CTCF (Cell Signaling Technology, catalog number 3418) and histone marks H3K27ac (Abcam, ab4729) and H3K27Me3 (Abcam ab6002). Independent biological replicates of each cell line (MCF10A, MCF10AT1, MCF10CA1a) were performed as described, including the optional step of snap freezing of chromatin and excluding the optional third washing step (104). Additionally, the chromatin was precleared with Pierce protein A/G beads for 2 hours at 4°C prior to incubation with antibodies. For CTCF we used 20ul of antibody (Cell Signaling Technologies, 3418S) and 150ug of chromatin for each sample. For H3K27ac 10ug of chromatin was used and 2ug of antibody, while for H3K27me3 15ug of chromatin was used with 4ug of antibody. Two biological replicates were generated per cell line from independent cultures and library preparations.

### ATAC-Seq library preparation

The OMNI-ATACseq protocol was followed as previously described, with an optimized 5 minutes of nuclear extraction rather than 3 minutes (105,106). Two biological replicates were generated per cell line from independent cultures and library preparations.

### Micro-C processing

Micro-C data from each technical replicate (library) was processed from raw reads into contact maps using guidelines outlined by Dovetail Genomics (107). Paired reads were aligned to the hg38 human genome assembly (NCBI GRCh38) using BWA mem (version 0.7.17; settings: -5SP -T0) (108). Pairtools (version 0.3.0) was then used to parse ligations, sort reads, and remove PCR duplicates using the parse (settings: --min-mapq 40 --walks-policy 5unique --max-inter-align-gap 30), sort, and dedup (settings: --mark-dups) commands (109). Alignment files were generated using pairtools split to create .pairs and .sam files, and samtools view (version 1.17; settings: -bS) to create .bam files (110). Final .bam alignment files were sorted and indexed using samtools sort and index commands. The .bam files were used to extract QC metrics using a script from Dovetail Genomics, and calculate complexity using preseq lc_extrap (settings: - bam -pe -extrap 2.1e9 -step 1e7 -seg_len 1000000000). Pairs files were used to generate contact maps using juicer_tools pre (version 1.22.01), and normalized using SCALE (111).

For merged contact maps, we first merged pairs files using pairtools merge, and then ran juicer_tools pre command on the resulting .pairs files. In total, we generated contact maps for each library (technical replicates), each sample (biological replicates), each cell type, and we created one fully merged map including all reads for a precise map of shared features. Micro-C maps for individual technical replicates and cell types are available on GEO (accession GSE254045).

### Micro-C copy number variation analysis

Structural variant and copy number variations were detected from contact maps using NeoLoopFinder calculate-cnv at 1Mb resolution (-e uniform; **Supp. File 2**) (112). These values were averaged over the course of large-scale variations and used as correction factors for differential loop analysis (see **Differential loop, TAD, gene, and peak analysis**, below).

### Regions excluded in analysis

Denylists of regions to ignore for feature calling were generated based on regions where SCALE normalization factors were unable to be calculated at 5kb or 10kb, ignoring single bins and merging continuous areas within 100 kb. Activity-by-contact analysis also combined this denylist with the ENCODE denylist for hg38 (113).

### Micro-C compartment calling

Compartments were called using CALDER (version 2.0; R version 4.2.1) at 10kb resolution and SCALE normalization (114). We also used FAN-C (version 0.9.21) to calculate eigenvectors at 100kb using SCALE normalization to build saddle plots, and oriented eigenvectors manually based on overlap with active genes (115). A table of subcompartments for each cell type is available on GEO (accession GSE254045).

### Micro-C topologically associated domain calling

Topologically associated domains (TADs) were called using SpectralTAD (version 1.18.0) at 10kb resolution with SCALE normalization, a window size of 300, and a level of 3, with a quality filter applied (116). We then created a unified TAD list by merging TADs that overlap at both ends between cell types, with a maximum gap of 10kb (1 pixel). We then combined this analysis with continuous insulation score (IS) calculations from FAN-C insulation command at 10kb resolution with SCALE normalization. We then used the FAN-C boundaries command to detect IS boundaries and only kept TADs that also overlapped with an IS boundary. Tables of TADs and boundaries are included in **Supp. File 4 and 5**. Insulation score tracks for each cell type are available on GEO (accession GSE254045).

### Micro-C chromatin loop calling

Chromatin loops were called using SIP (version 1.6.2), run at 5kb and 10kb and then merged to 10kb (-g 2 -fdr 0.05). Loops were called in each cell type map using SCALE normalization, in addition to the combined map, and then merged similarly to TADs to create a unified loop list (**Supp. File 3**) (117). Cell type loops are available on GEO (accession GSE254045).

### RNA-Seq processing

All RNA-seq data was analyzed using the nf-core/rnaseq pipeline (118). Adapter and quality trimming was performed with Trim Galore (119). Reads were aligned to the hg38 reference genome using STAR (120) and gene expression was quantified using Salmon (121). Differentially expressed genes were called using DESeq2 (122). An adjusted *p*-value of 0.05 and a log2fold change of 1 were used as thresholds to select significant differential expression. A table of all differentially expressed genes and their DESeq2 metrics are listed in **Supp. File 6**. A full table of expression data for all genes can be found on GEO (accession GSE320216).

### ChIP-Seq processing and peak calling

ChIP-Seq data for CTCF, H3K27ac, and H3K27me3 was processed as detailed previously (41,123). In summary, adapters were cut (cutadapt v1.11) and low quality reads trimmed (Galaxy FASTQ Quality Trimmer 1.0.0; window 10, step 1, minimum quality 20). Reads were mapped to the human genome (hg38 canonical) using STAR version 2.4 (120) with splicing disabled (–alignIntronMax 1). Enriched regions (narrowPeak calls for CTCF and H3K27me3, broadPeak calls for H3K27ac) for each replicate were generated using MACS2 (Feng et al., 2012) and replicates were then evaluated using deepTools (124) to correlate alignments and IDR (125) to evaluate peak call reproducibility. After pooling replicates, MACS2 (126) was used to call peaks at high stringency (P-value <10e-5), these peaks were further filtered according to IDR cutoffs. ChIP-seq data is available on GEO (accession GSE98551 for CTCF, and GSE229295 for H3K27ac and H3K27me3).

### ATAC-Seq processing and peak calling

Read trimming and quality filtering was performed using FastQC (127) and TrimGalore (119). The filtered fastq were then downsampled to approximately 50 million reads per replicate. Reads were aligned to the hg38 reference genome using Bowtie2 (128). Mitochondrial, Multi-mapped, and low quality reads, and duplicates were removed using samtools (110). MACS2 (126) was used to call narrowPeaks, followed by IDR (125) to generate sample level peak sets. ATAC-seq data is available on GEO (accession GSE320215).

### Enhancer and promoter definitions

Gene promoters were defined as regions between 2000 bp upstream and 500 bp downstream of gene transcription start sites.

Potential enhancer regions were identified based on regions that contained both an ATAC-Seq and H3K27ac ChIP-Seq peak. For activity-by-contact analysis, potential enhancers were defined as 150,000 ATAC-Seq peaks with the highest levels of H3K27ac signal, but were subset for regions with H3K27ac peaks after running (see below).

### Compartmental saddle plots

Saddle plots were made manually in R. We selected three chromosomes that had no major karyotypic differences between cell lines and had high correlation of eigenvectors between replicates with the same signage (chr2, chr12, chr17). For each chromosome and cell type, we sorted eigenvectors into 20 bins. We then calculated the mean observed/expected values (using SCALE normalization) between regions belonging to different bins, and plotted it as a heatmap.

### Differential loop, TAD, gene, and peak analysis

Differential genes were calculated using DESeq2 (version 1.42.0) (122). Each cell type had 3 replicates, and a design of ∼cellType was used. No fold-change cutoff was applied. Genes with an adjusted p-value below 0.01 were considered significant.

Differential H3K27ac within ATAC-Seq peaks were calculated using a similar design, but with a p-value cutoff of 0.05.

Differential CTCF peaks were called using the CSAW package following the workflow described in the original publication (41), and an adjusted p-value cutoff of 0.1.

Differential loops were also identified using DESeq2, but with additional adjustments. Raw, un-normalized loop counts were pulled from each technical replicate map, resulting in 8 values per cell type (4 technical replicates for each of 2 biological replicates per cell type). An LRT design was used of ∼technicalRep + biologicalRep + cellType, with ∼technicalRep + biologicalRep as the comparison. Size factors were provided manually based on NeoLoopFinder output (**Supp. File 2**). Regions of duplication were identified based on NeoLoopFinder CNV output and spectral karyotypes. For each duplicated region, the CNVs across the region were averaged and used as custom size factors to correct for differences in chromosomal copy number between cell lines. Differential loops were identified based on a fold-change cutoff of 1.5 and an adjusted p-value cutoff of 0.025.

Differential TAD boundaries were detected using an alternative method. Insulation scores were pulled from all TAD boundaries at the technical replicate level (8 values per cell type). A T-test was then applied for each pairwise comparison of cell types, and p-values were adjusted using FDR. Boundaries with an adjusted p-value below 0.01 were considered significant.

Differential loops, TAD boundaries, and genes were clustered based on their patterns of change across the three cell types using k-means clustering of centered and scaled normalized counts.

### Integration of Cancer Genome Atlas data

RNA-Seq data from the breast cancer (BRCA) cohort of the Cancer Genome Atlas was used to analyze whether genes differentially expressed in the MCF10 progression model are also differentially expressed in breast cancer tumor tissues (n=1102) from patients comparted to normal control tissues (n = 113). Normalized gene transcript counts from RSEM were downloaded from Xena (129). Protein coding genes that are differentially expressed between tumors and paired peritumors in TCGA-BRCA data were identified using GEPIA3 using DESeq2 with a log2 fold-change cutoff of 1 and an adjusted p-value cutoff of 0.05. A total of 5,289 differential genes were identified from the TCGA-BRCA data: 3,168 with significantly higher expression in tumor samples, and 2,121 with significantly higher expression in normal samples (130). Overall survival curves for select genes were generated using GEPIA (130).

### Integration of additional breast cancer structural data

Chromatin loops and TAD boundaries identified in the MCF10 progression model were also examined in other triple-negative breast cancer cell lines and patient samples using data from GEO (accession GSE167150) (49). Insulation score tracks were calculated as for MCF10 cell lines, using FAN-C at 10kb resolution and KR normalization. Chromatin loop counts were extracted using KR normalization at 10kb resolution.

### Micro-C feature overlap and aggregate analysis

Micro-C feature overlap and analysis was conducted in R primarily using the *GenomicRanges*, *InteractionSet*, and *mariner* packages (131–133). To overlap chromatin loops with other features such as promoters and enhancers, we used loop anchors at 10kb resolution and allowed features within 10kb (±1 Hi-C pixel) of the loop anchors to be considered overlapping. We used this broad definition for loop overlap to account for the fact that loops are non-punctate and show increased contact frequency within several bins on either side of the called loop pixel, as evidenced by both individual loci and aggregate loop analysis. Aggregate matrices of loop pixels and TAD boundaries were generated using *mariner* and visualized using *plotgardener* (133,134).

### Activity-by-contact

The activity-by-contact was used based on Fulco et al. 2019 with slight adjustments (135). To allow for direct comparison of all enhancer-promoter pairs across cell lines, we manually defined potential enhancer regions and used the same input for each cell type. These potential enhancer regions were defined as they typically are in the pipeline, by finding 150,000 ATAC-Seq peaks with the highest H3K27ac levels. The output was filtered using a suggested ABC score cutoff of 0.025 to identify likely enhancer-promoter pairs. To allow for direct comparison with our other enhancer analysis, we filtered the output based on the enhancer regions also overlapping H3K27ac, which still represented a majority of the valid pairs identified.

### Matched enhancer-promoter sets

Covariate-matched subset selection among non-looped enhancer-promoter pairs was performed using the *matchRanges* function from the *nullranges* package (136,137). Enhancer-promoter pair distance or mean Micro-C contact frequency were used as covariates. Matching was done with the stratified matching method without replacement.

### Gene ontology and gene set enrichment analysis

Gene ontology (GO) term enrichment was performed in R using the *gprofiler2* package (137). Gene set enrichment analysis (GSEA) was performed with the *GSEA* software, using size factor normalized RNA-seq counts as input (138) and the Hallmark H1 gene set.

### ATAC-Seq motif analysis

Motif analysis of ATAC-Seq peaks within strengthened and weakened loop anchors was conducted using the HOMER suite findMotifsGenome.pl script with size given(139). ATAC-Seq peaks within the anchors of static loop anchors were used as background.

### H3K27ac peak pileup analysis

H3K27ac ChIP-Seq analysis in the anchors of gained, lost, and static loops was conducted using deeptools (124). Alignment files were normalized using RPGC with the bamCoverage function, and adjusted using scale factors generated from edgeR TMM normalization factors of counts from overlapping H3K27ac and ATAC-Seq peaks (140). Aggregate profile plots were then created using the plotProfile command.

### Genomic data visualization

Micro-C contact frequency maps, aggregate analysis plots, gene annotations, and genomic signal tracks (RNA-Seq, ChIP-Seq, ATAC-Seq) were visualized and plotted in R using the *plotgardener* package (134).

## ACKNOWLEDGEMENTS

We thank Kathleen Quinn and Joseph Boyd (University of Vermont) for their work building the ChIP-Seq libraries and processing the ChIP-Seq data, respectively. We thank Darawalee Wangsa and Danny Wangsa in the CCR Genetics Branch and OMICS Technology Facility at the NCI for their expert SKY analysis. We thank Alquassem Abuorquob (University of Vermont) for his assistance in ChIP-Seq library preparation. We thank Jordan Zhang, Misha Gattengo, and Sierra Wilson at Dovetail Genomics for coordinating Micro-C library preparation and preliminary analysis. STR analyses and Next Generation Sequencing was in part done with the assistance of the Vermont Integrated Genomics Resource (VIGR) at the Vermont Cancer Center, University of Vermont and the Genomics Sequencing Facility (GSF) at Greehey Children’s Cancer Research Institute UT Health San Antonio. The results published here are in part based upon data generated by the TCGA Research Network: https://www.cancer.gov/tcga. We thank the members of the Misteli and Stein labs for feedback.

## DECLARATIONS

## Ethics approval and consent to participate

Not applicable.

## Consent for publication

Not applicable.

## Availability of data and materials

The Micro-C, ATAC-Seq, and RNA-Seq datasets generated and analyzed during the current study are available on GEO (SuperSeries GSE320319; SubSeries GSE254045, GSE320215, GSE320216). CTCF ChIP-Seq data was previously published under GEO accession GSE98551. H3K27ac and H3K27me3 ChIP-Seq data was previously published under GEO accession GSE229295.

The code used to generate figures from the current study are available on Github at the following repository: https://github.com/ksmetz/MCF10-MicroC.

## Competing interests

The authors declare they have no competing interests.

## Funding

This work was supported in part by the Intramural Research Program of the National Institutes of Health (NIH), National Cancer Institute, Center for Cancer Research (grant no. ZIABC010309-24 to T.M.), the Northern New England Clinical Translation Network (grant no. GM115516 to G.S.), funding from the National Cancer Institute (grant no. P01CA240685 to G.S., J.S., and S.F.; grant no. 1F32CA220935-01A1 to A.F.), and funding from the Charlotte Perelman and Arthur Jason Perelman Fund for Cancer Research (G.S. and J.S.). The contributions of the NIH author(s) were made as part of their official duties as NIH federal employees, are in compliance with agency policy requirements, and are considered Works of the United States Government. However, the findings and conclusions presented in this paper are those of the author(s) and do not necessarily reflect the views of the NIH or the U.S. Department of Health and Human Services

## Authors’ contributions

KSMR, TM, AF, HG, JS, and GS conceptualized the project, designed the experiments, and interpreted the results. TM, SF, GS, and JS obtained funding. KSMR and AF executed experiments. KSMR and HG performed computational processing and analysis. SF conducted ChIP-Seq processing. KHH coordinated SKY karyotyping. KSMR and TM drafted the manuscript. KSMR, AF, HG, KHH, JS, GS, and TM participated in reviewing and editing the manuscript.

## SUPPLEMENTARY FILES

**Supplementary File 1:** TableS1-microcMetrics.xls

Supplementary File 1. Micro-C library quality metrics for individual technical replicates, biological replicates, and cell types.

**Supplementary File 2:** TableS2-CNVs.xls

Supplementary File 2. CNVs from NeoLoopFinder (100kb resolution). Columns indicate bin coordinates (A-C), and CNV values for MCF10A (D), MCF10AT1 (E), and MCF10CA1a (F).

**Supplementary File 3:** TableS3-loops.xls

Supplementary File 3. Chromatin loop summary table for total loop set (n = 29,205). Columns include loop anchor coordinates (A-J), loop call status by cell type (K-N), SIP AP score (O), loop name (P), loop span length (Q), average and maximum un-normalized counts (R-S), log2 fold-change (T-V), adjusted p-value (W), differential status by pairwise comparison (X-Z), un-normalized counts by technical replicate (AA-AX), maximum log2 fold-change (AY), pairwise comparison with greatest fold-change (AZ), differential cluster (BA), Z-score normalized counts (BB-BE), and variance stabilized counts (BF-BH).

**Supplementary File 4:** TableS4-TADbounds.xls

Supplementary File 4. TAD boundary summary table for total boundary set (n = 17,097). Columns include boundary coordinates (A-C), insulation scores by technical replicate (D-AA), adjusted p-values, difference in insulation score, and differential status for each pairwise comparison (AB-AJ), differential status across all comparisons (AK), differential cluster (AL), average insulation score per cell type (AM-AO), and Z-score normalized insulation scores (AP-AS).

**Supplementary File 5:** TableS5-TADs.xls

Supplementary File 5. TAD summary table for total set of TADs (n = 13,231). Columns include TAD boundary coordinates (A-J), and whether the TAD was called in each cell type (K-M).

**Supplementary File 6:** TableS6-differentialGenes.xls

Supplementary File 6. Differentially expressed genes from the MCF10 progression series (n = 8,840). Columns indicate gene coordinates (A-E), gene identifiers (F-I), base mean expression (J), pairwise log2 fold-change (K-M), adjusted p-values (N-P), and differential status (Q-S), differential cluster (T), un-normalized counts by replicate (U-AC), Z-score normalized counts (AD-AF), and variance stabilized counts (AG-AI).

## SUPPLEMENTAL FIGURE LEGENDS

**Figure 1 - Figure Supplement 1.**
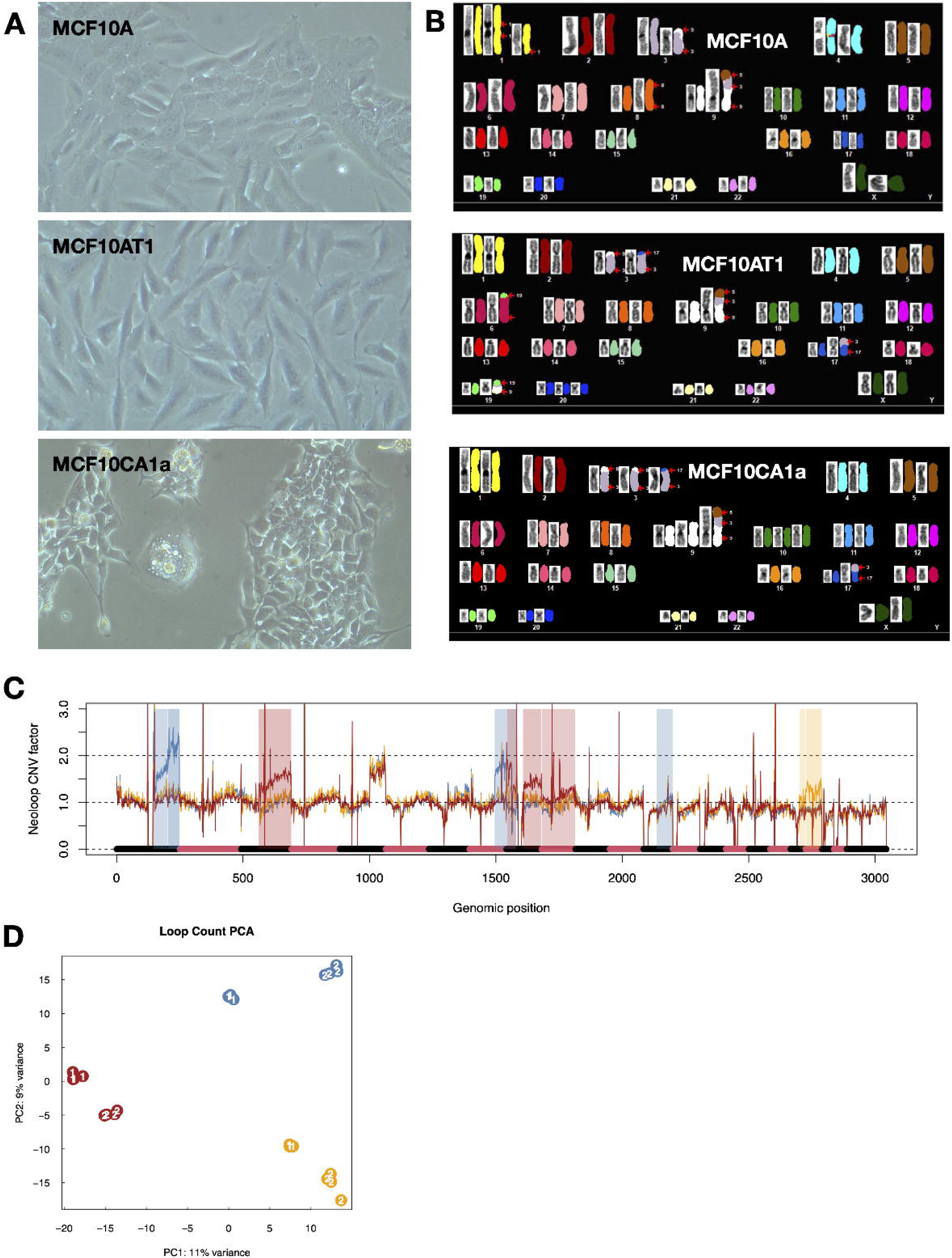
Karyotypic analysis and Micro-C loop reproducibility. **(A)** Brightfield microscopy images of MCF10A, MCF10AT1, and MCF10CA1a cell lines. **(B)** Representative SKY karyotype example images from MCF10A, MCF10AT1, and MCF10CA1a cell lines. **(C)** Copy number variation (CNV) factors for loops across the genome as generated from NeoloopFinder. Highlighted regions indicate areas of karyotypic differences that were corrected for in the identification of differential loops; blue is regions with higher CNV in MCF10A, yellow is regions with higher CNV in MCF10AT1, and red is regions with higher CNV in MCF10CA1a. Regions that have shared karyotypic abnormalities among all three cell lines, such as the q arm of chromosome 5 which is duplicated and translocated to chromosome 9 in all cell types, did not require correction. **(D)** Micro-C chromatin loop count principal component analysis (PCA). Blue circles indicate MCF10A samples, yellow indicate MCF10AT1, and red indicates MCF10CA1a.

**Figure 1 - Figure Supplement 2.**
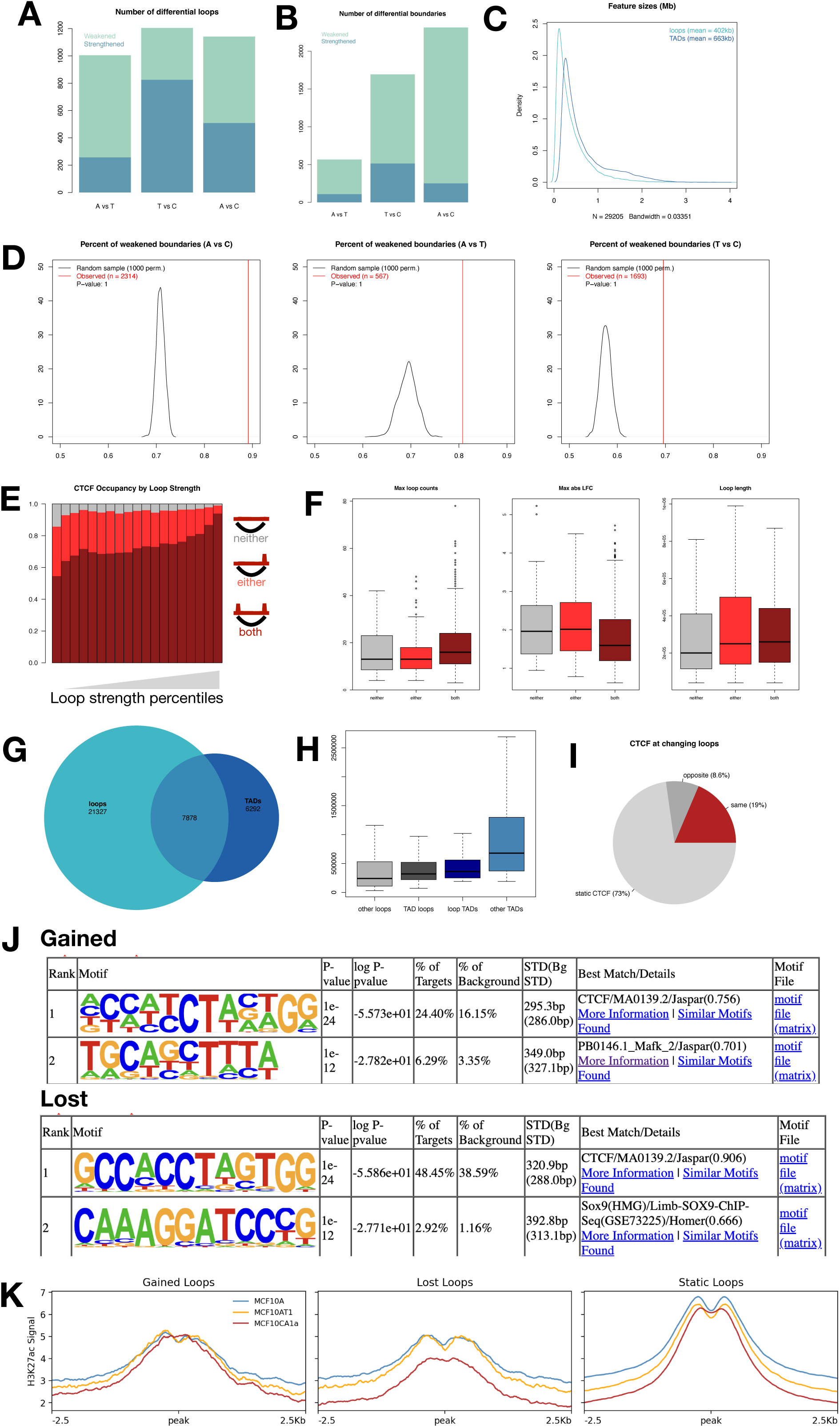
Differential loop and TAD feature details. **(A)** The number of chromatin loops that exhibit a significant increase (blue) or decrease (green) in contact frequency between each pairwise comparison of cell types. **(B)** The number of topologically associating domain (TAD) boundaries that exhibit a significant increase (blue) or decrease (green) in insulation score between each pairwise comparison of cell types. **(C)** Size distributions of chromatin loops and TADs. **(D)** Permutation test results of weakened boundaries observed in each pairwise comparison as compared to a random sampling. **(E)** Percent of loops with one or both anchors overlapping CTCF ChIP-Seq peaks based on the strength of the loop (20 bins). **(F)** Boxplots of (left to right) maximum loop Micro-C counts, maximum loop count log2 fold-change, and loop length based on loop-CTCF class; neither (grey), one anchor overlap (light red), or both anchors overlap (dark red). **(G)** Venn diagram showing the degree of overlap between chromatin loops and TADs. **(H)** Feature sizes of loops that do not overlap TADs (light grey), loops that do overlap TADs (dark grey), TADs that do overlap loops (dark blue), and TADs that do not overlap loops (light blue). **(I)** Pie chart of the percentage of differential loops that have a correlated change in CTCF binding at either anchor; light grey indicates static CTCF peaks, dark grey indicates CTCF peaks that change in the opposite way as the loop (i.e. a strengthened loop with loss of CTCF binding), and dark red indicates CTCF peaks that change in the same way as the loop (i.e. a strengthened loop with gain of CTCF binding). **(J)** De novo motif results from HOMER for ATAC-Seq peaks within the anchors of gained/strengthened (top) and lost/weakened (bottom) chromatin loops. **(K)** H3K27ac ChIP-Seq aggregate profiles at the anchors of gained/strengthened, lost/weakened, and static loops.

**Figure 1 - Figure Supplement 3.**
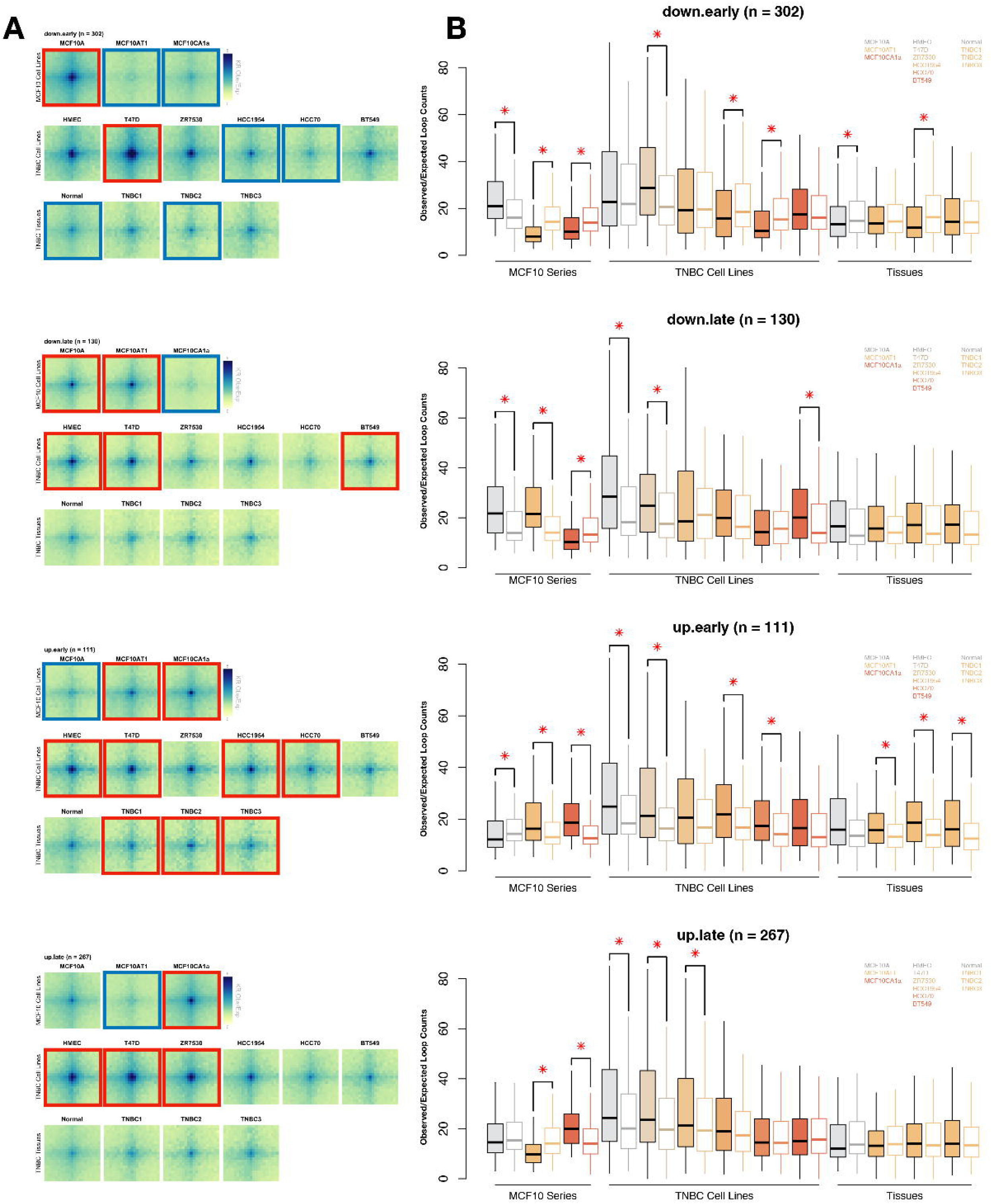
Differential MCF10 loops are conserved in cancer cell lines. **(A)** Aggregate loop analysis of chromatin loops from each differential cluster as found in the MCF10 series compared to TNBC cell lines and primary patient tissues (Kim, Han & Chun et al. 2022). Red boxes indicate loops that have significantly higher observed/expected contact frequency among loops in that cluster as compared to a random sampling of an equal number of chromatin loops that are static in the MCF10 series, as determined in (B). **(B)** Boxplots representing quantifications of the aggregate plots shown in (A). Filled boxes indicate the counts from loops in each differential cluster, while empty boxes indicate a matched sample of static loops that are of similar size. Stars indicate cell lines where there is a significant difference between the counts of the given loops and the static matched set (p <= 0.05, Wilcoxon rank sum test).

**Figure 1 - Figure Supplement 4.**
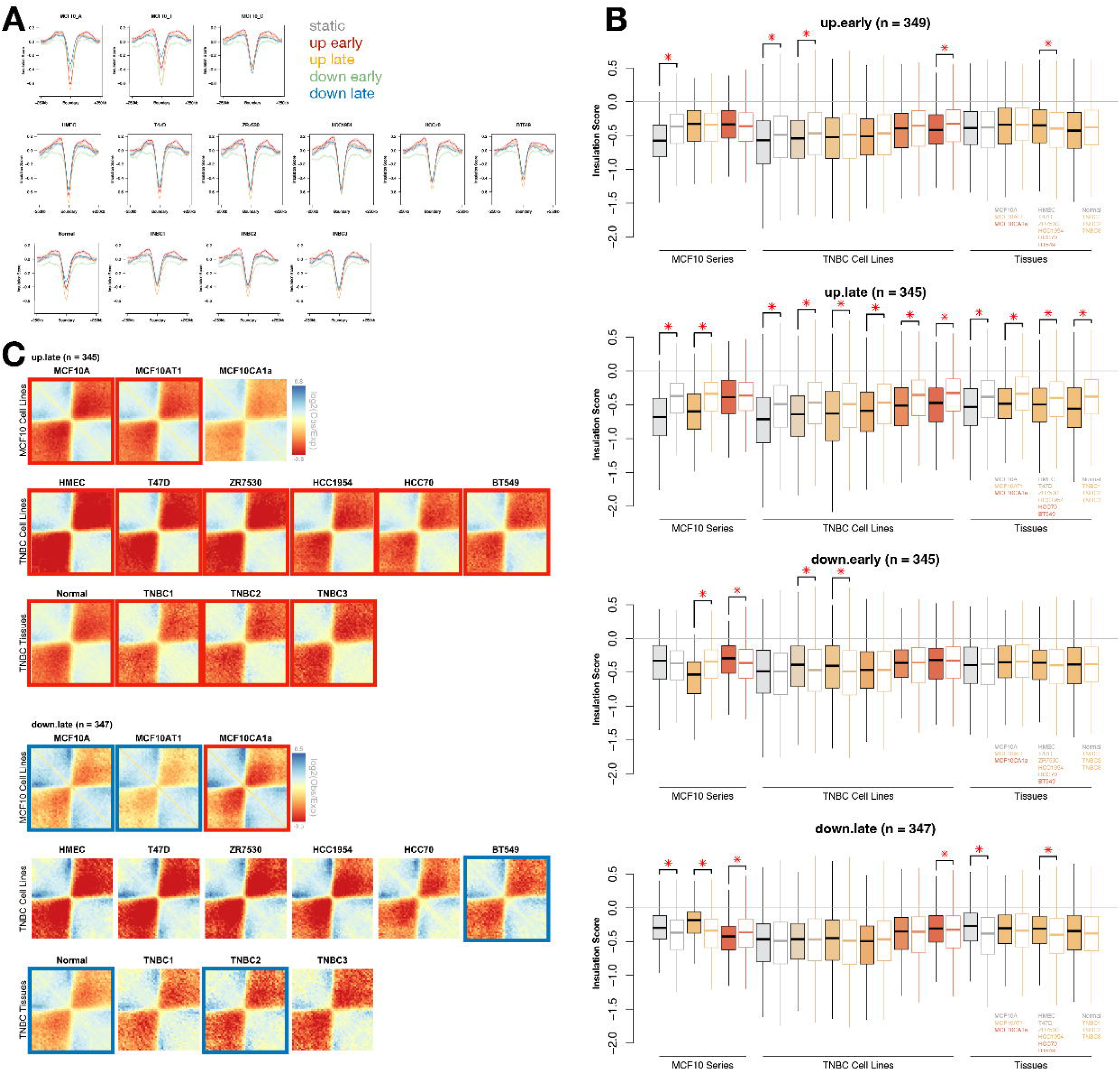
Differential TAD boundaries are conserved in cancer cell lines and tissues. **(A)** Average insulation score profile of TAD boundaries based on their differential cluster in MCF10 progression and in TNBC cell lines and primary patient tissues (Kim, Han & Chun et al. 2022). **(B)** Boxplots showing the distribution of insulation scores at MCF10 boundaries of various differential clusters. Filled boxes indicate the insulation scores from TADs in each differential cluster, while empty boxes indicate the distribution of scores at the same number of static TAD boundaries. Stars indicate cell lines where there is a significant difference between the insulation score of boundaries and the static matched set (p <= 0.05, Wilcoxon rank sum test). **(C)** Aggregate plots showing the log2 of observed/expected counts at MCF10 boundaries in MCF10 cell lines as well as cell lines from Kim, Han & Chun et al. 2022.

**Figure 2 - Figure Supplement 1.**
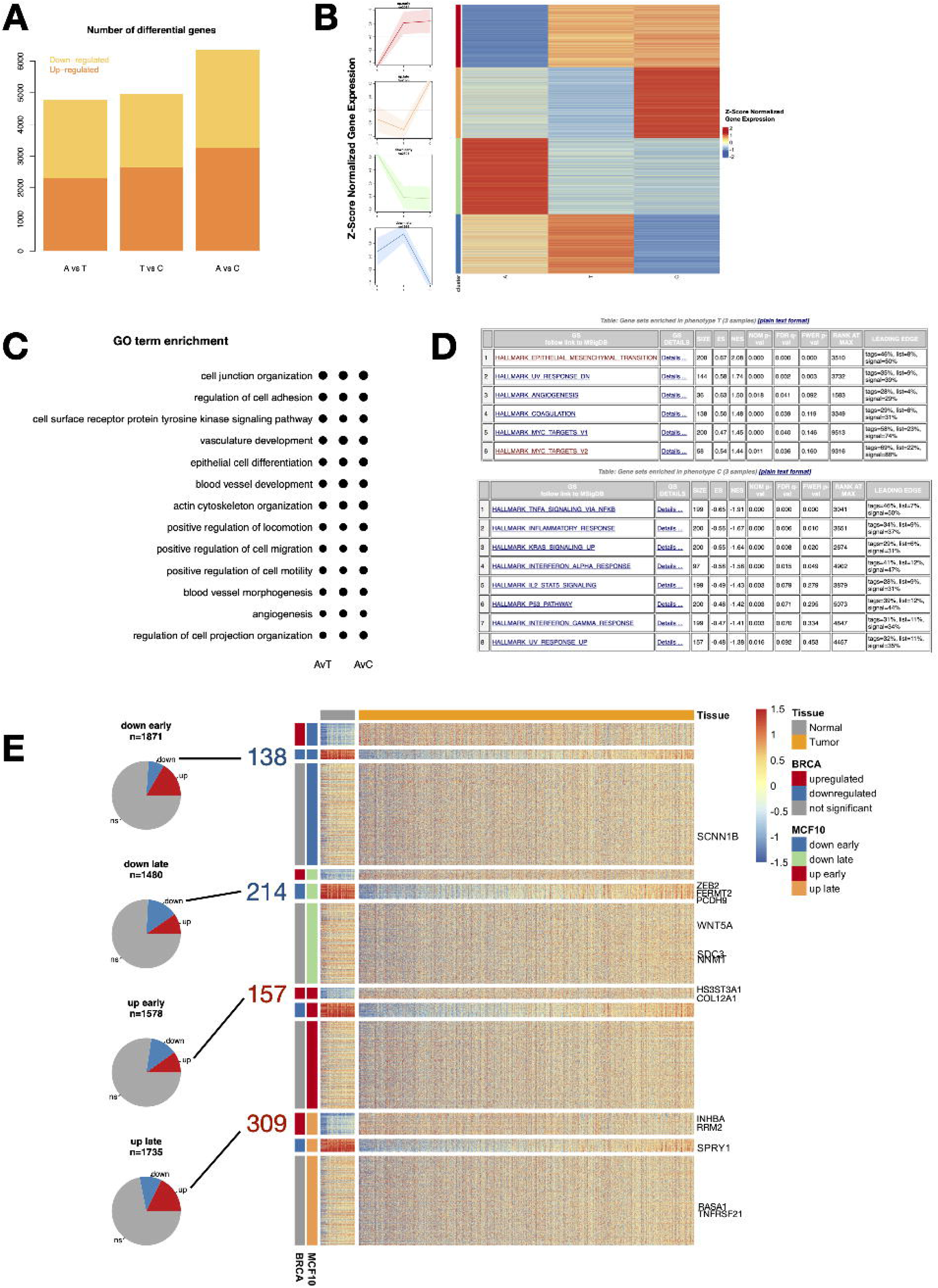
Differential gene expression patterns across breast cancer progression includes cancer-relevant pathways. **(A)** The number of genes that exhibit a significant increase (orange) or decrease (gold) in expression between each pairwise comparison of cell types. **(B)** Differential genes clustered by their timing of change, depicted in line plots and heatmap. **(C)** Gene ontology (GO) term enrichment for genes differential expressed in each pairwise comparison of cells. **(D)** Gene set enrichment analysis (GSEA) results showing gene sets enriched among genes differentially expressed early (MCF10AT1, top) or late (MCF10CA1a, bottom) in breast cancer progression. **(E)** A heatmap shows the expression of all differential genes in the MCF10 progression series among normal (grey) and tumor (gold) tissue samples from the Cancer Genome Atlas breast cancer cohort. Genes are clustered first by their pattern of change in MCF10, then their differential status in the patient data. Example genes used in the study are listed on the right. Each cluster also shows a pie chart on the left depicting the percent of genes that are shared in the patient data, with genes that change in the same direction highlighted.

**Figure 2 - Figure Supplement 2.**
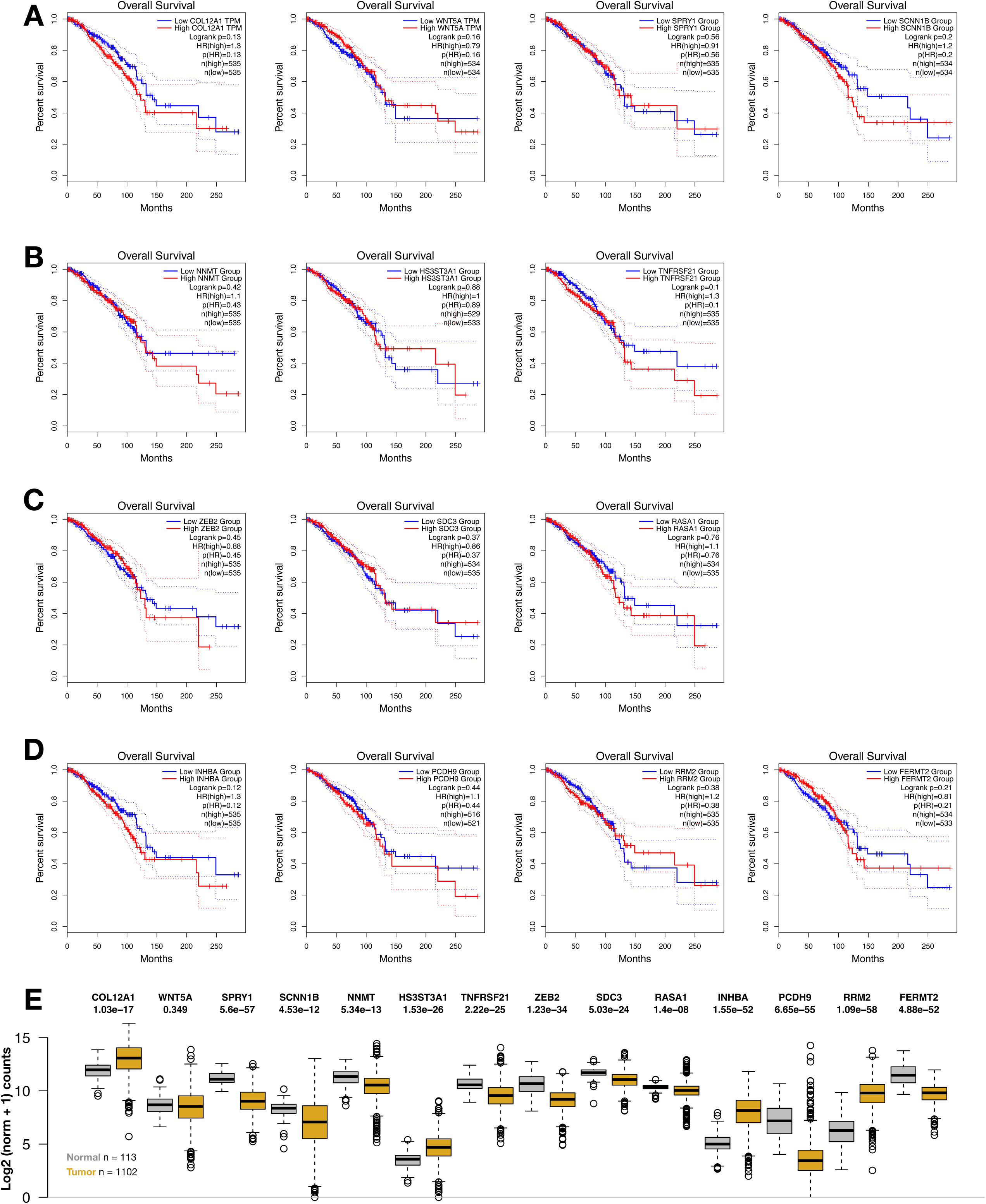
Clinical relevance of select genes misregulated in MCF10. (A-D) Overall survival curves from the Cancer Genome Atlas for fourteen select genes highlighted in the main figures (A), Supplemental Figure S7 (B-C), and Supplemental Figure S12 (D). **(E)** Gene expression data from the Cancer Genome Atlas for fourteen select genes. Grey boxplots represent expression values from normal samples and gold represent tumor samples. P-values represent the Wilcoxon rank-sum test comparing the normal and tumor expression levels for each gene.

**Figure 2 - Figure Supplement 3.**
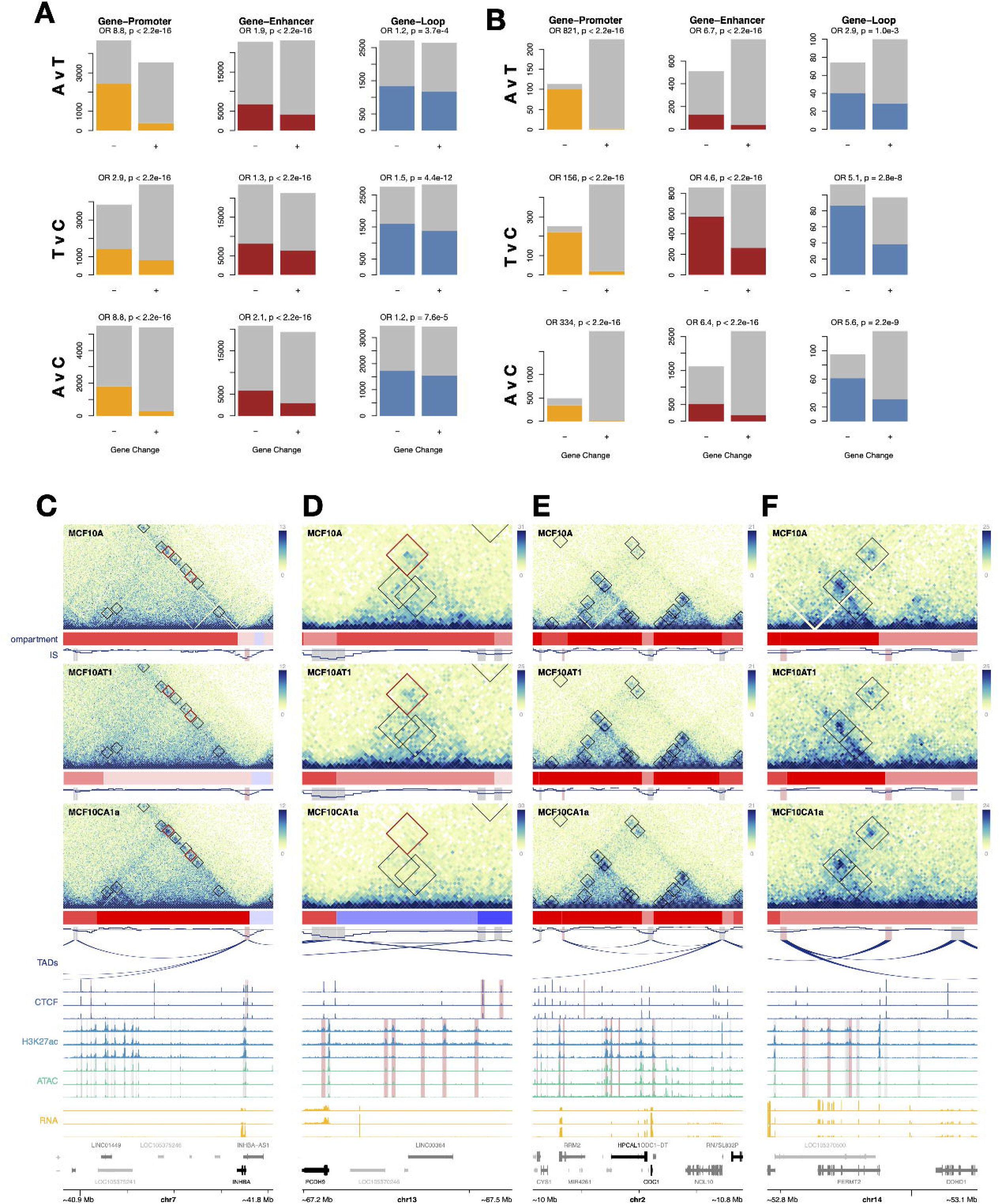
Differentially expressed genes overlap differential and static loops with differential distal regulatory regions. **(A)** The number of promoter H3K27ac peaks (left), distal enhancer H3K27ac peaks (middle), or loops (right) that change in a positive (grey) or negative (gold, red, blue) direction based on their overlap with up-regulated or down-regulated genes. **(B)** Same plots as (**E**) but subset for significantly differential features. **(C-F)** Examples of genes that are differentially regulated in the same direction in MCF10 progression and patient samples and which overlap with chromatin loops in MCF10 cell lines. Black boxes show loop annotations, while red boxes indicate differential loops. Red compartment tracks indicate A compartments, while blue indicates B compartments. In CTCF signal tracks, red highlights indicate differential CTCF peaks. In H3K27ac and ATAC-Seq signal tracks, red highlights indicate differential enhancers as determined by changes in H3K27ac. Genes highlighted in black are differentially expressed.

**Figure 2 - Figure Supplement 4.**
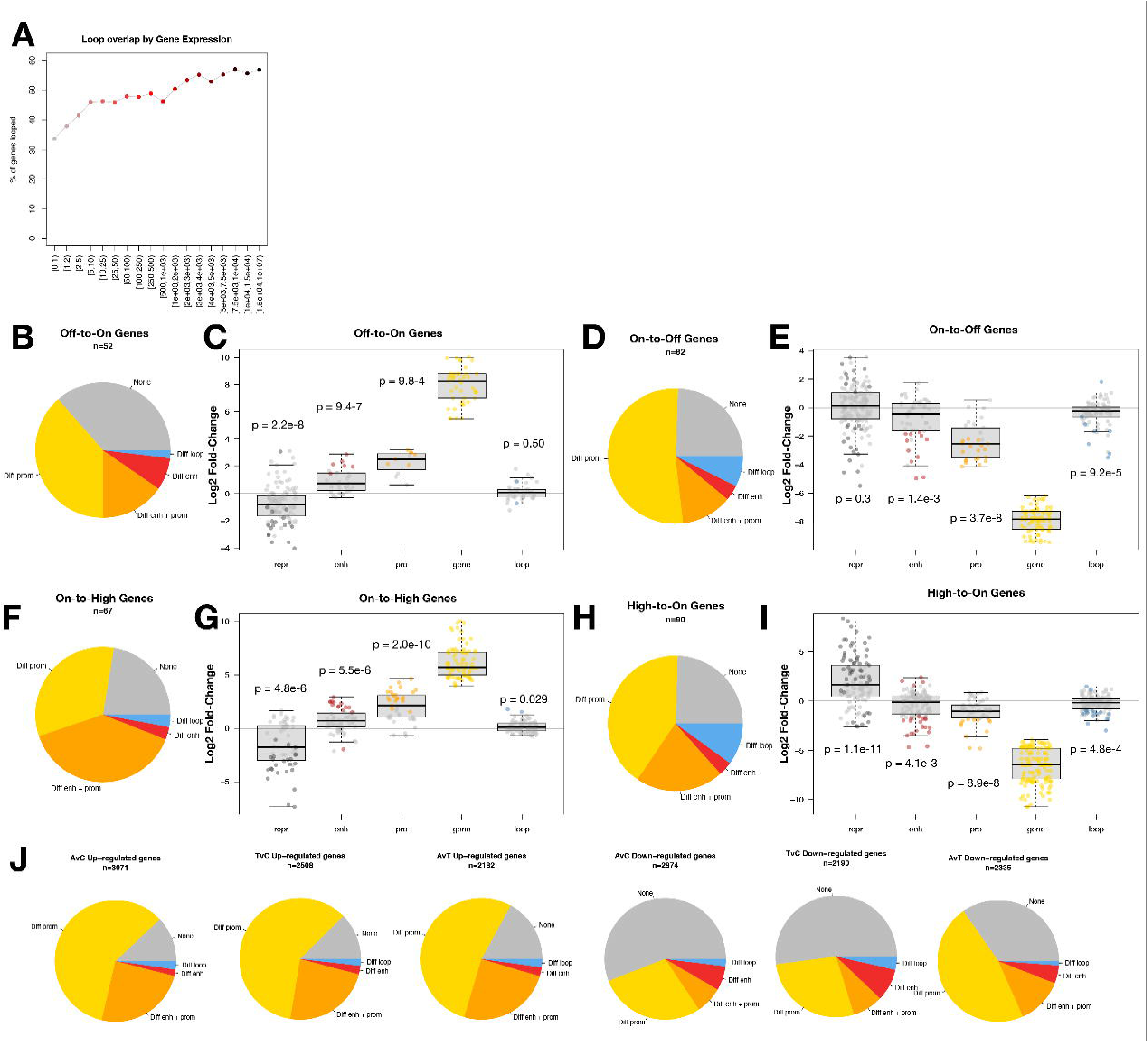
Relationship between differential gene expression, chromatin loops and distal enhancers is consistent based on expression levels and cancer stages. **(A)** Percentage of genes that overlap with loop anchors based on gene expression level. **(B)** Percentages of off-to-on genes that have gained H3K27ac at promoters, distal enhancers, both, or gained loops. **(C)** Log2 fold-change of distal H3K27me3 (grey), distal H3K27ac (red), promoter H3K27ac (orange), gene expression (yellow), and loop strength (blue), when overlapped. Grey dots indicate features that do not change significantly, while colored points are significantly differential features. P-values represent Wilcoxon tests comparing the means of each class to 0. Non-significant (n.s.) represents p-values above 0.01. **(D-I)** Same as (B-C), but for on-to-off genes (D-E), on-to-high genes (F-G), and high-to-on genes (H-I). **(J)** Percentages of up-regulated genes (left) and down-regulated genes (right) that have correlated changes in H3K27ac at promoters (gold), distal enhancers (red), both (orange), or loop contact frequency (blue), by pairwise comparison of cell types.

**Figure 2 - Figure Supplement 5.**
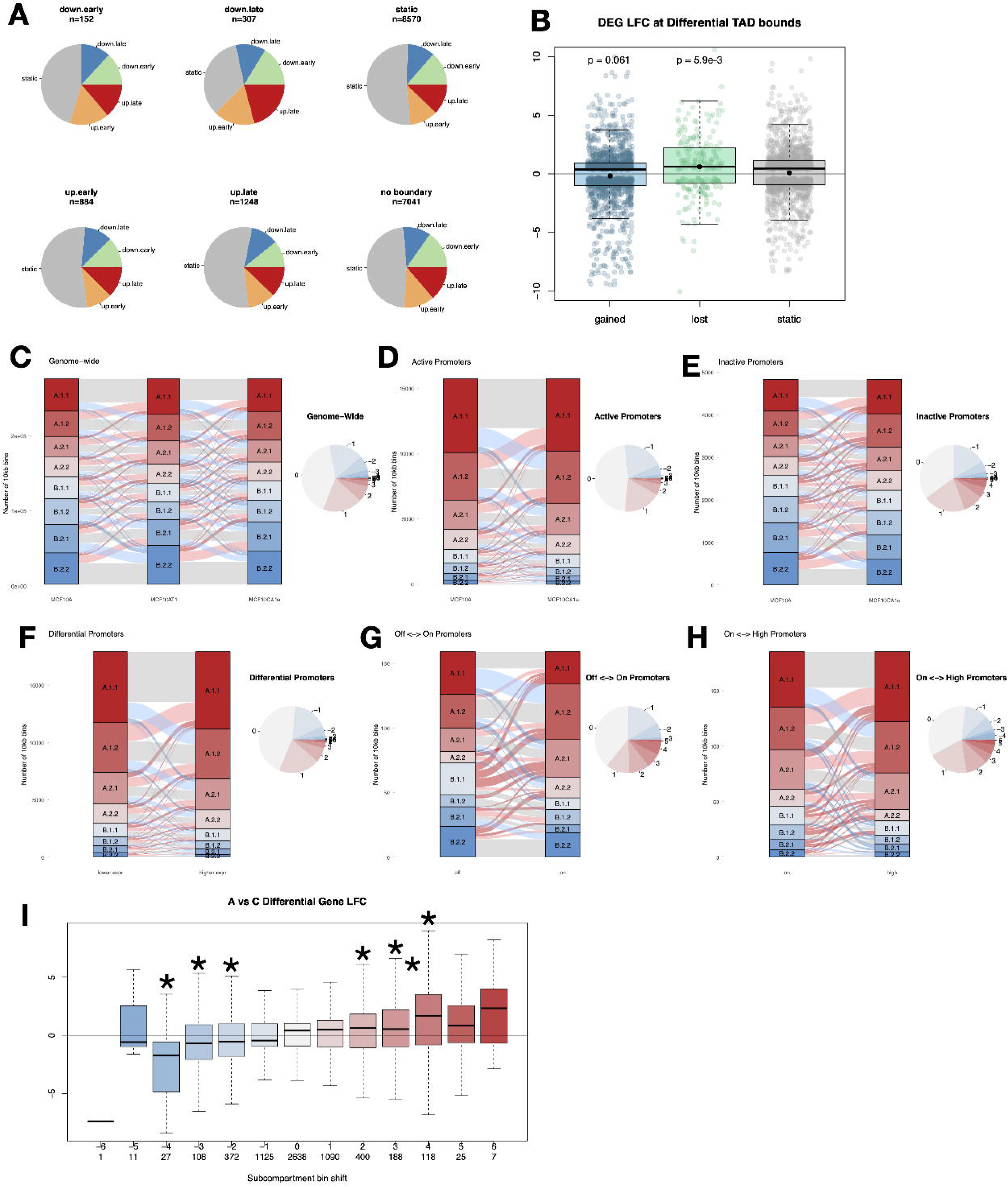
Differential TAD boundaries and subcompartment shifts have a subtle relationship to gene expression. **(A)** Pie charts showing the differential status of genes within 50kb of TAD boundaries of various differential clusters. **(B)** Boxplots of the log2 fold-change of genes at the boundaries of TAD boundaries that are either weakened (blue), strengthened (green), or static (grey). P-values represent Wilcoxon rank sum tests comparing the mean of each set to the static set. **(C)** Alluvial plots showing the transition of subcompartments from MCF10A to MCF10AT1, and from MCF10AT1 to MCF10CA1a. Grey ribbons indicate bins that have the same subcompartments between cell types, while red ribbons shift more A-like and blue ribbons shift more B-like. Darker ribbons shift by more than one subcompartment. A pie chart to the right summarizes the proportion of shifts based on the number of subcompartments they change by. **(D-E)** Alluvial plot showing the difference in subcompartments for the promoters of (D) actively expressed or (E) non-expressed genes that have similar gene expression in MCF10A and MCF10CA1a. Plots and pie charts are colored as in (C). **(F-H)** Alluvial plots showing the difference in subcompartments for the promoters of genes that are differentially expressed between any two cell types (F), change from on to off (G), or change from on to high (H). For each figure, the left bar shows the subcompartments in the cell line with lower expression and the right bar is from the cell line with higher expression. The pie charts summarize the shifts as in (C). **(I)** A boxplot indicating the distribution of gene log2 fold-change between MCF10A and MCF10CA1a for genes with promoters in bins that shift by various numbers of subcompartments. Negative numbers (blue) indicate bins that shift to more B-like subcompartments, while positive numbers (red) indicate shifts to more A-like subcompartments. The numbers below indicate the number of bins that fit into each category. Stars indicate significant differences in mean gene log2 fold-change compared to genes that do not change subcompartments between the two cell types (grey), as indicated by a p-value of 0.01 or less (Wilcoxon rank sum test).

**Figure 3 - Figure Supplement 1.**
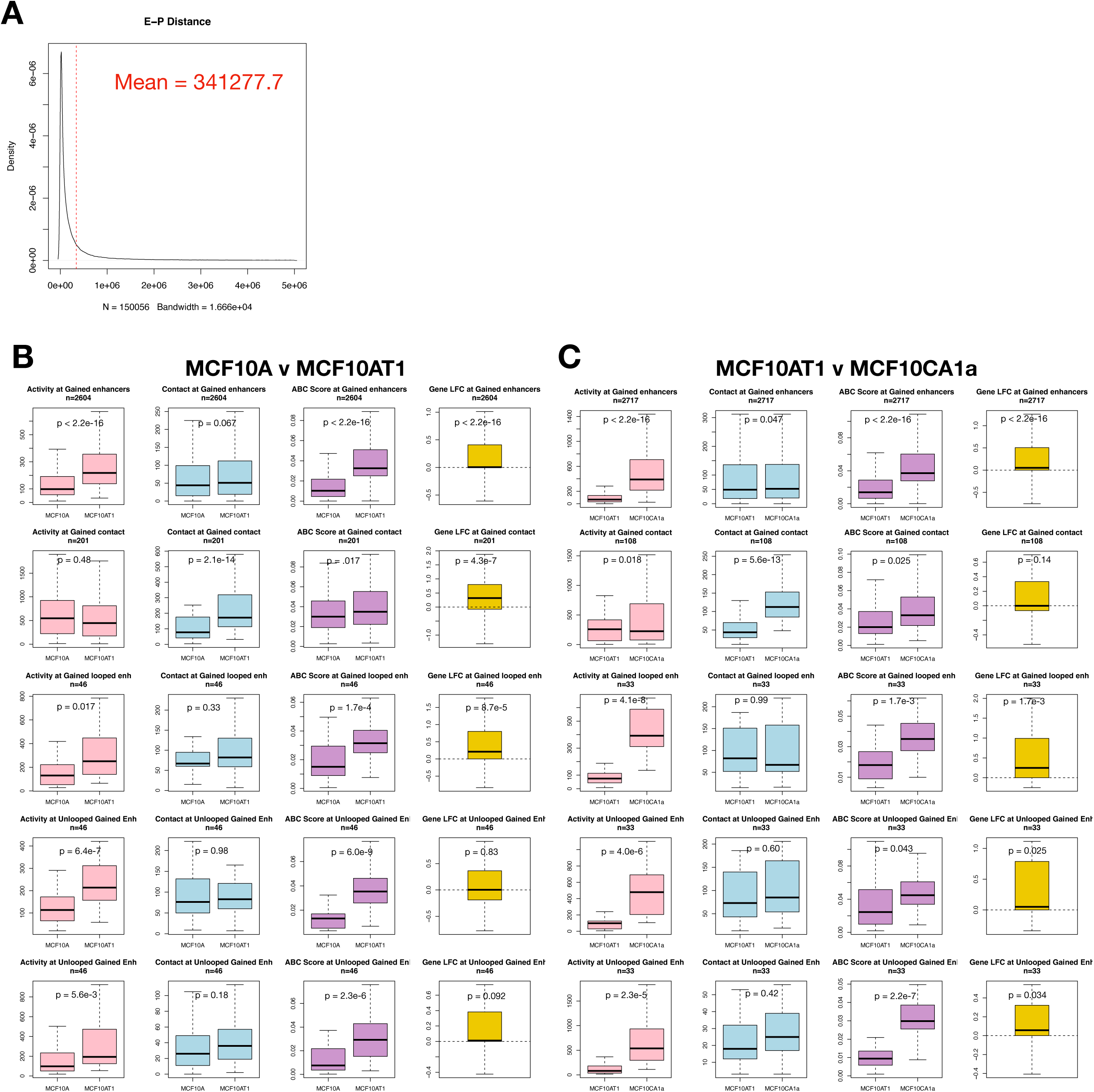
Enhancer-promoter pair details. **(A)** Distribution of enhancer-promoter distances as predicted by the activity-by-contact (ABC) model. **(B)** Boxplots of distal enhancer H3K27ac (pink), enhancer-promoter contact (blue), and ABC score (purple), as well as gene log2 fold-change (yellow) for enhancer promoter pairs that feature (top to bottom) differential H3K27ac among enhancers, differential enhancer-promoter contact frequency, differential H3K27ac for looped enhancer-promoter pairs, contact-matched non-looped enhancer-promoter pairs, and distance-matched non-looped enhancer-promoter pairs. Comparison shows changes between MCF10A and MCF10AT1. **(C)** Same as (B) but showing comparisons between MCF10AT1 and MCF10Ca1a.

**Figure 4 - Figure Supplement 1.**
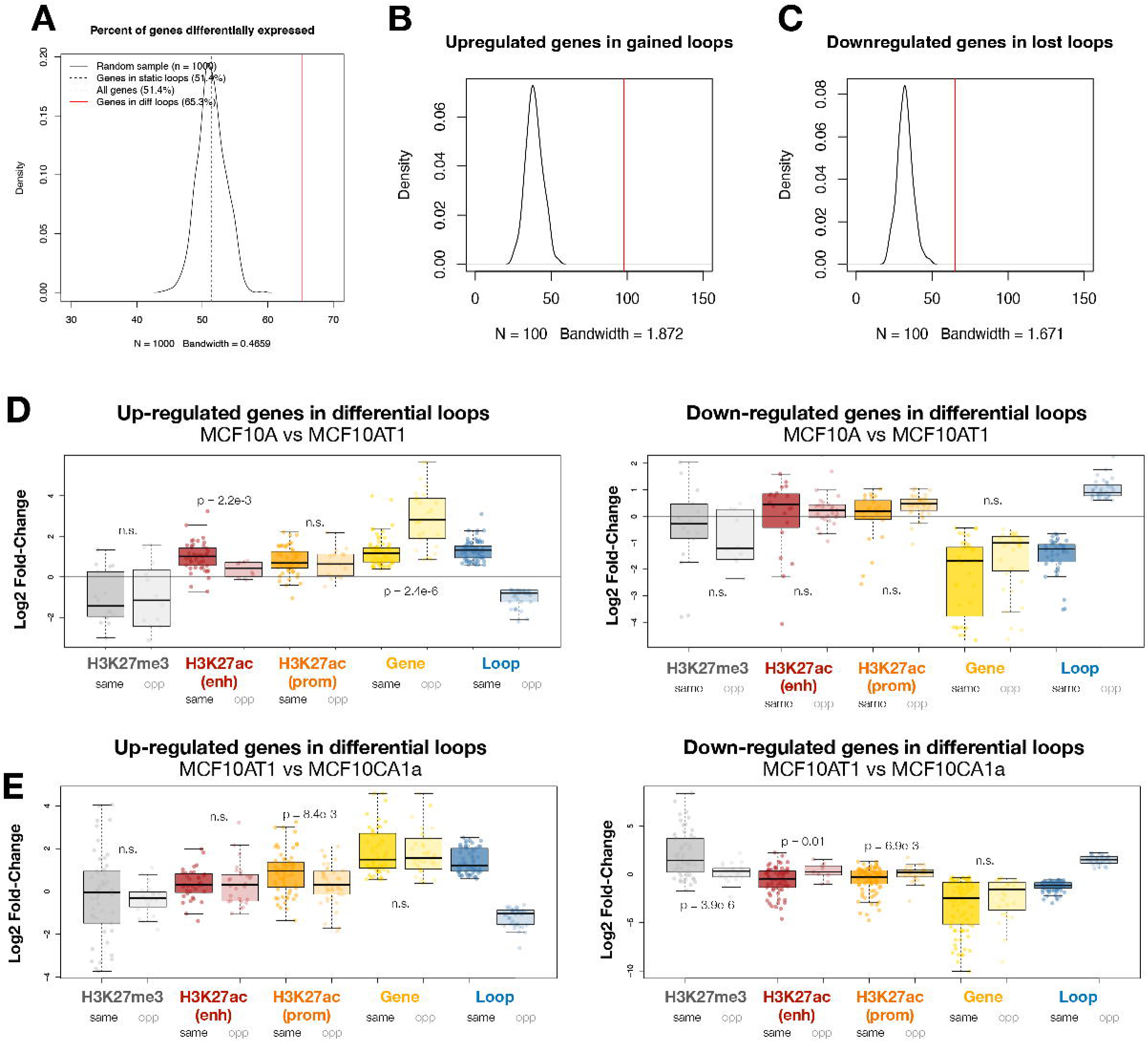
Differential genes within differential loops. **(A)** The number of differential expressed genes among all genes (dashed light grey line), genes overlapping static loop anchors (dashed dark grey line), genes overlapping differential loop anchors (red line), and a permutation test of a random sampling of a similar number of genes (black line). **(B)** Permutation test results (n=1,000) for the number of up-regulated genes that overlap with strengthened/gained loops (red) compared to a random sampling (black). **(C)** Permutation test results (n=1,000) for the number of down-regulated genes that overlap with weakened/lost loops (red) compared to a random sampling (black). Boxplots in **(D)** and **(E)** represent log2 fold-change between MCF10A and MCF10AT1 (D) or MCF10AT1 and MCF10CA1a (E) of distal H3K27me3 (grey), distal H3K27ac (red), promoter H3K27ac (orange), gene expression (yellow), and loops (blue) among upregulated or downregulated genes that overlap with gained loops (darker colors) or lost loops (lighter colors. Boxplots are defined as in (A). P-values represent T-tests comparing the mean values of the features at loops that change in the same and those that change in opposite directions from the differential genes at their anchors. Non-significant (n.s.) p-values are any p- values above 0.01.

**Figure 4 - Figure Supplement 2.**
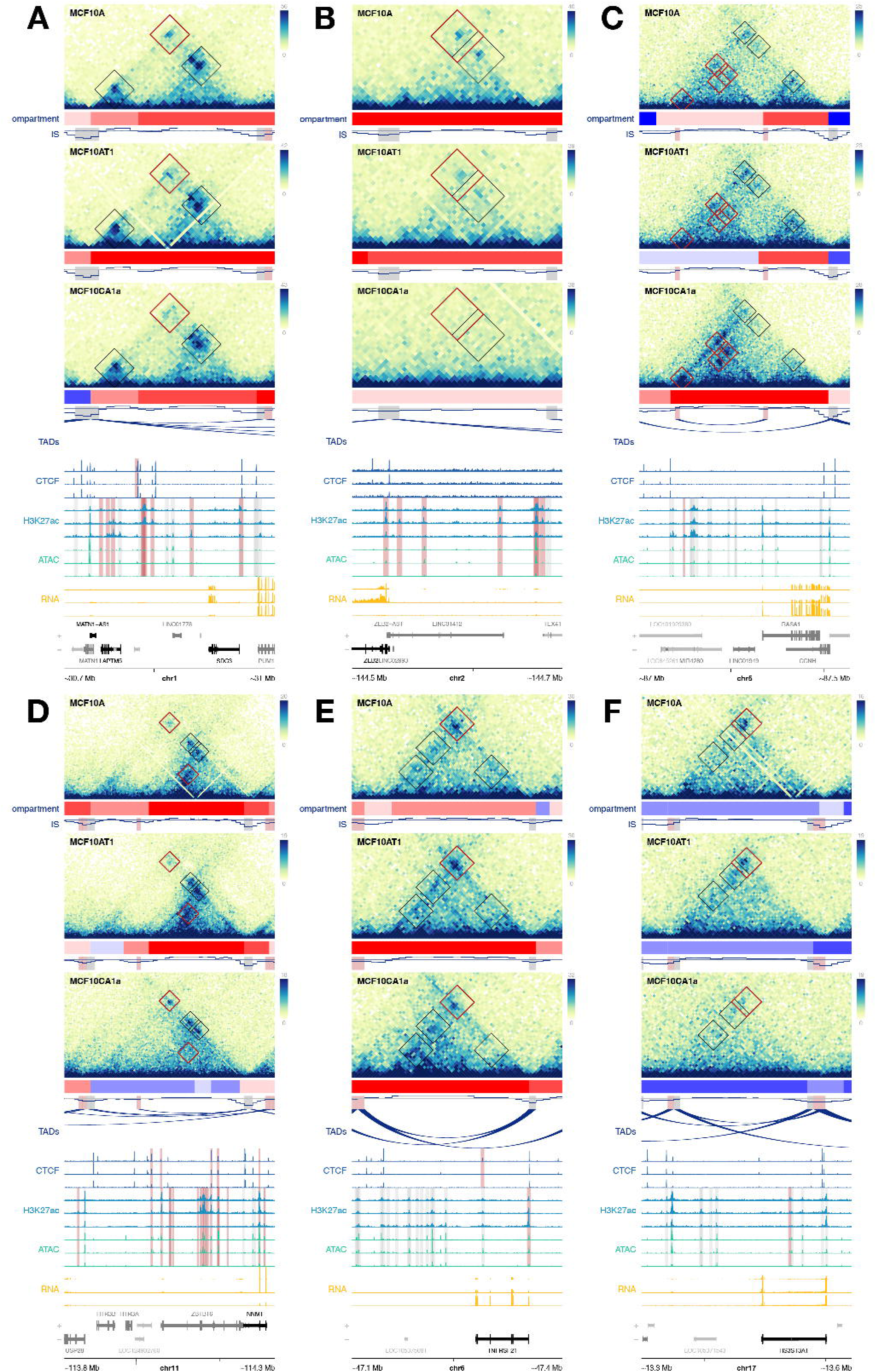
Examples of differentially expressed genes at differential loop anchors. (A-C) Examples of genes that are differentially expressed in MCF10 progression and overlap with anchors of differential genes that change in the same direction. Black boxes show loop annotations, while red boxes indicate differential loops. Red compartment tracks indicate A compartments, while blue indicates B compartments. In CTCF signal tracks, red highlights indicate differential CTCF peaks. In H3K27ac and ATAC-Seq signal tracks, red highlights indicate differential enhancers as determined by changes in H3K27ac. Genes highlighted in black are differentially expressed. **(D-F)** Examples of genes that are differentially regulated in MCF10 progression and overlap with anchors of differential genes that change in the opposite direction. Plots are annotated as in (A-C).

## Notes

https://www.ncbi.nlm.nih.gov/geo/query/acc.cgi?acc=GSE320319

### Competing Interest Statement

The authors have declared no competing interest.

### Summary of Updates

Fixed figure order (supplemental figures at end)

